# A Novel *Tmem119-tdTomato* Reporter Mouse Model for Studying Microglia in the Central Nervous System

**DOI:** 10.1101/665893

**Authors:** Chunsheng Ruan, Linlin Sun, Alexandra Kroshilina, Lien Beckers, Philip L. De Jager, Elizabeth M. Bradshaw, Samuel Hasson, Guang Yang, Wassim Elyaman

## Abstract

Microglia are resident immune cells of the central nervous system (CNS). The exact role of microglia in the physiopathology of CNS disorders is not clear due to lack of tools to discriminate between CNS resident and infiltrated innate immune cells. Here, we present a novel reporter mouse model targeting a microglia-specific marker (TMEM119) for studying the function of microglia in health and disease. By placing a reporter cassette (GSG-3xFlag-P2A-tdTomato) between the coding sequence of exon 2 and 3’UTR of the *Tmem119* gene using CRISPR/Cas9 technology, we generated a *Tmem119-tdTomato* knock-in mouse strain. Gene expression assay showed no difference of endogenous *Tmem119* mRNA level in the CNS of *Tmem119*^tdTomato/+^ relative to control Wild-type mice. The cells expressing tdTomato-were recognized by immunofluorescence staining using commercially available anti-TMEM119 antibodies. Using immunofluorescence and flow cytometry techniques, tdTomato^+^ cells were detected throughout the CNS, but not in peripheral tissues of adult *Tmem119*^tdTomato/+^ mice. In addition, aging does not seem to influence TMEM119 expression as tdTomato^+^ cells were detectable in the CNS of older mice (300 and 540 days old). Further immunofluorescence characterization shows that the tdTomato^+^ cells were highly colocalized with Iba1^+^ cells (microglia and macrophages) in the brain, but not with NeuN- (neurons), GFAP- (astrocytes) or Olig2- (oligodendrocytes) labeled cells. Moreover, flow cytometry analysis of brain tissues of adult mice demonstrates that the majority of microglial CD45^low^CD11b^+^ cells (96.6%) are tdTomato positive. Functionally, using a laser-induced injury model, we measured time-lapse activation of tdTomato-labeled microglia by transcranial two-photon microscopy in live *Tmem119*^tdTomato/+^ mice. Taken together, the *Tmem119-tdTomato* reporter mouse model will serve as a valuable tool to specifically study the role of microglia in health and disease.

## 1. Introduction

Microglia constitute the resident macrophages of the central nervous system (CNS)(Graeber and Streit, 1990), with a panoply of functions including immune surveillance (Hanisch and Kettenmann, 2007), maintenance of neuronal networks (Wu et al., 2015), and injury repair (Colonna and Butovsky, 2017). Microglia are phagocytic cells, and clearance of debris is thought to be a key function, especially in aging (Pluvinage et al., 2019). They can also secrete a wide range of cytokines including chemokines. Historically, microglia were often described as resting (i.e., ramified), but this phrasing failed to convey the dynamic remodeling of their fine processes and constitutive immunosurveillance activity, and are now termed homeostatic (Boche et al., 2013; Lawson et al., 1993). This morphology was in contrast to a more amoeboid phenotype microglia can also obtain, which was referred to as macrophages, but now is more commonly referred to as activated to avoid confusion with infiltrating cells (Arcuri et al., 2017). Both morphological and physiological properties have been used to identify microglial cells. The markers frequently used to identify microglia are CD68, MHC class II, CD11b or Iba1. However, since these markers are also expressed on other myeloid cells, they fail to distinguish between resident microglia and infiltrating macrophages.

The importance of microglia in most neurodegenerative diseases, such as multiple sclerosis and Alzheimer’s disease, is increasingly appreciated (Pena-Altamira et al., 2016; Wang and Colonna, 2019). As microglia are highly plastic, the local disease environment, where microglia are in close interaction with other CNS cell types, differs between different pathologies and different regions leading to various phenotypes. Therefore, the microglial phenotype is disease-dependent, influenced by neighboring cell types and disease pathologies such as aggregated amyloid-beta (Bachiller et al., 2018). Neurodegenerative diseases such as multiple sclerosis are associated with infiltration of peripheral monocytes/macrophages that contribute to disease pathogenesis (Meyer-Luehmann and Prinz, 2015). Many markers used for the identification of microglia are also present in macrophages, as both cell types are myeloid and share many transcriptional programs. Given a lack of reliable markers that can distinguish between resident microglia and infiltrating myeloid cells, the specific function of microglia in CNS diseases has been difficult to ascertain.

During the past few years, our group and others have generated bulk and single-cell transcriptomics data to identify specific markers of human and mouse microglia (Bachiller et al., 2018; Bennett et al., 2016; Butovsky et al., 2014; Chiu et al., 2013; Fahrenhold et al., 2018; Gosselin et al., 2017; Hickman et al., 2013; Katsumoto et al., 2014; Konishi et al., 2017; Marta Olah, 2019; Olah et al., 2018). Bennett *et al* identified Transmembrane protein 119 (TMEM119) as a specific marker of human and mouse microglia and generated an anti-TMEM119 monoclonal antibody to detect microglia *in situ* (Bennett et al., 2016). TMEM119, an evolutionarily conserved 58-kDa type 1 transmembrane protein, was originally identified as a regulator of osteoblast differentiation. It is also expressed on human osteoblasts and follicular dendritic cells. Mice homozygous for a targeted deletion of the *Tmem119* gene exhibit growth retardation associated with delayed endochondral bone ossification and impaired osteoblast differentiation(Hisa et al., 2011). Although the role of TMEM119 in bone development is becoming clear, whether TMEM119 is involved in microglia development and function is not known.

In an effort to develop a reliable tool for studying microglia, we have generated a novel microglia reporter mouse strain where the tdTomato fluorescence gene was knocked-in between the coding sequence of exon 2 and the termination codon in the 3’ UTR of the *Tmem119* gene thus preserving endogenous *Tmem119* expression. Using immunofluorescence and flow cytometry analyses, we demonstrate that the TMEM119-tdTomato signal is detected in the brain and spinal cord, is specific to microglia and is not detected in peripheral myeloid cells. Using two-photon microscopy, we also show that our *Tmem119-tdTomato* knock-in mouse strain is reliable for live imaging of microglia in naïve mice as well as in mice that were exposed to brain injury.

## 2. Materials and Methods

### 2.1. Mice

*Tmem119-tdTomato* reporter mice were generated in BioCytogen Co, Ltd (Beijing, China). Wild-type C57BL/6J mice were purchased from the Jackson Laboratory (Maine, USA) and were crossed with the *Tmem119-tdTomato* mice. Animals were housed in the pathogen-free animal facility at Columbia University Medical Center, in accordance with the guidelines of the Committee of Animal Research at Columbia University and the National Institutes of Health animal research guidelines as set forth in the Guide for the Care and Use of Laboratory Animals. All studies were performed in compliance with procedures approved by Columbia University Institutional Animal Care and Use Committee.

### 2.2. Generation of the Tmem119-tdTomato knock-in mice using CRISPR/Cas9

The Tmem119-tdTomato knock-in mouse strain was generated using CRISPR/Cas9 technology. Briefly, eight single-guide RNAs (sgRNA, see **supplementary File 1**) were designed to target the sequence of the stop codon of murine *Tmem119* locus. The activity of the CRISPR/Cas9 was assessed using a UCA kit. The results of genotyping with the UCA kit showed that one sgRNA (sgRNA #4) was transcribed successfully with required concentration (**supplementary File 1)**. Following embryo microinjection using a micromanipulator, embryos were cultured to the two-cell stage, followed by transfer into pseudopregnant C57BL/6 females. The founders were genetically examined by amplifying sequences spanning the 5’ and 3’ junction and including the entire inserted transgene. The detailed workflow of the mouse line design, plasmid construction, Southern blot study design, microinjection and founder genotyping is shown (**Supplementary File 1**).

### 2.3. DNA isolation and genotyping

Mouse genomic DNA was isolated from tail biopsies (0.5-1 cm) and following overnight digestion at 55◻ into 500 uL of buffer containing 100 mM Tris-HCl (pH 8.0), 5 mM EDTA (pH 8.0), 200 mM NaCl, 0.2% SDS and 0.1 mg/mL Proteinase K. The samples were gently mixed and cooled down at room temperature for 10-15 min. After centrifugation at 12000 rpm for 15 min, 400 uL of the supernatants were collected and gently mixed with the same volume of isopropanol. After centrifugation at 12000 rpm for 10 min, the pellets were washed by gently mixing with 700 uL of 75% ice-cold ethanol. After centrifugation at 12000 rpm for 5 min, the DNA pellets were harvested and air-dried at room temperature for 5 min. The DNA was dissolved in 100 uL of distilled water at 55◻ for two hours, and 100 ng DNA were used for PCR reaction. Wild-type primers (562bp): Forward: 5’-CAGAACCTCCGGTCTCCAGCTAGAG-3’; Reverse: 5’-AGAGAAGTGGTGCGTTAGGGTGAAG-3’; Knock-in primers (299bp): Forward: 5’-CCACCACCTGTTCCTGTACG-3’; Reverse: 5’-AGAGAAGTGGTGCGTTAGGGTGAAG-3’. The PCR conditions were as follows: 1) 95◻ for 15 min; 2) 35 cycles at 94◻ for 45 sec, 60◻ for 1 min, and 72◻ for 1 min; 3) 72◻ for 5 min. PCR products were separated on 2% Agarose gel. Detailed analysis of the genotyping strategy is provided (**Supplementary File 2)**.

### 2.4. RNA isolation and real-time quantitative PCR

Animals were humanely killed by CO2 and transcardially perfused with ice-cold PBS. The left-side half brains were harvested and frozen in dry ice and stored at −80C. RNA was isolated using RNeasy Plus Micro Kit (#74034; QIAGEN, USA), and 10 uL of RNA was reversely transcribed into cDNA using the High-Capacity cDNA Reverse Transcription Kit (#4368814; Thermo Fisher Scientific, USA) following the manufacturer’s instructions. The *Tmem119* gene expression was detected by real-time quantitative PCR using TaqMan Fast Advanced Master Mix (#4444557; Thermo Fisher Scientific) and *Tmem119* TaqMan Gene Expression Assay (#Mm00525305_m1; Thermo Fisher Scientific). *Gapdh* TaqMan Gene Expression Assay (#Mm99999915_g1; Thermo Fisher Scientific) was applied as an internal amplification control. The PCR conditions were as follows: 50◻ for 2 min, 95◻ for 2 sec, and followed by 40 cycles at 95◻ for 1 sec and 60◻ for 20 sec. The gene expression of *Tmem119* relative to *Gadph* was quantified by the ΔΔCt method.

### 2.5. Immunofluorescence staining

Animals were humanely killed by CO2 and and transcardially perfused with ice-cold PBS. The right-side half brains were harvested and fixed in 4% paraformaldehyde (made in PBS; #15710, Electron Microscopy Sciences, USA) at 4◻ for 6 days. After dehydration in 30% sucrose (made in PBS; #57-50-1, Affymetrix, USA), 30-μm thick free-floating sections were sliced using the Cryostat CM3050S (Leica, Germany). For immunostaining, sections were washed in PBS three times (10 min each), and incubated in blocking buffer (3% BSA (#A7906, Sigma, USA) in PBS supplemented with 0.1% Triton X-100 (#9002-93-1; Sigma)) at room temperature for one hour. Primary antibodies (rabbit anti-TMEM119 (1:500;#ab209064; Abcam, USA), mouse anti-TMEM119 (1:100; #853302, BioLegend, USA), rabbit anti-Iba1 (1:500; #019-19741; Wako, Japan), mouse anti-NeuN (1:500; #MAB377, Millipore, USA), rabbit anti-GFAP (1:500; #ZO334; Dako, USA), rabbit anti-Olig2 (1:500;#AB9610; Millipore, USA) diluted in 1% BSA (in PBS) were further applied to the sections at 4°C overnight. After being washed in PBS four times (10 min each), the sections were incubated in species-matched secondary antibodies conjugated with 488 diluted in 1% BSA (in PBS) at room temperature for one hour. Sections were washed again in PBS for four times (10 min each), and mounted on slides with mounting media with DAPI (#NC9524612; Vector Laboratories, USA), and imaged using confocal microscope (LSM700, Zeiss, Germany).

### 2.6. Flow cytometry

Animals were humanely killed by CO2 and blood was collected followed by perfusion with ice-cold PBS. Brain, spinal cord, lung, heart, liver, spleen, kidney and intestine were harvested, and homogenized in 2% FBS (in PBS). The cells were collected by centrifuging at 300g, 4◻ for 10 min and were resuspended in MACS buffer (1% FBS in PBS supplemented with 0.4% 0.5 M EDTA). The red blood cells were lysed by incubating with the same volume of ACK lysis buffer (#A1049201, Gibco, USA). Cells were resuspended after being washed twice in MACS buffer and analyzed by BD Accuri C6 flow cytometer (BD Biosciences, USA). The blood samples were incubated with 10× volume of RBC Lysis Solution (#1045703, QIAGEN) at room temperature for 10 min. After being washed twice in MACS buffer, the cells were resuspended and analyzed by BD Accuri C6. Microglia were isolated as previously described (Butovsky et al., 2014; Olah et al., 2012; Verheijden et al., 2015). Briefly, mice were humanely killed by CO2 and transcardially perfused with ice-cold PBS. The brain tissues were quickly dissected and single cell suspensions were prepared and separated using a 37%/70% Percoll gradient (#GE17-0891-01, GE Healthcare). Myelin was removed from the top layer and mononuclear cells were isolated from the interface. The cells were incubated with mouse anti-CD45-FITC (1:200; #103108, BioLegend) and mouse anti-CD11b-APC (1:80; #101212, BioLegend) for 30 minutes on ice in the dark. Cell suspensions were analyzed using the BD FACS Canto (BD Biosciences), and data were analyzed with FlowJo software (TreeStar, USA).

### 2.7. Two-photon laser injury inside the cortex

A highly localized injury was achieved by focusing a two-photon laser beam (◻1 μm in size) in the superficial layer of the cortex through the open skull window. The wavelength of the two-photon laser was set at 800 nm and the laser power was ◻60–80 mW at the sample. The beam was parked at the desired position for approximately 3-5 seconds to create a small injury site as indicated by a bright autofluorescent sphere (◻30 μm in diameter) around the focal point of the beam. The injury was confined to the area ◻25–35 μm in diameter around the laser focal point, because microglia within this area lost their tdtomato immediately after laser ablation, whereas those ◻50 μm from the injury site still responded to the ablation. The laser-induced focal ablation is a useful injury model, as the site and degree of injury are easy to control, and the response of microglia toward the injury is highly reproducible.

### 2.8. *In vivo* imaging of microglia with two-photon microscopy

To investigate microglia in vivo in intact brain, tdTomato-labeled microglia were imaged by two-photon microscopy through a small craniotomy. Briefly, 2-3 month-old *Tmem119-tdTomato* mice were anesthetized intraperitoneally with ketamine (200 mg/kg body weight) and xylazine (30 mg/kg body weight) in 0.9% NaCl solution. A region (◻1 mm in diameter) over primary sensory cortex was first thinned with a high-speed drill under a dissecting microscope and then opened either with a needle or forceps. A drop (◻200 μl) of artificial mouse cerebrospinal fluid (ACSF) was applied on the exposed region for the duration of the experiment. The skull surrounding the open window was attached to a custom-made steel plate to reduce respiratory-induced movement. The animal was placed under a two-photon microscope (Scientifica Hyperscope, East Sussex, UK) while still under anesthesia and temperature maintained with a heating pad (~37°C). The two-photon laser was tuned to the excitation wavelength for tdTomato (1000 nm). All experiments were performed using a 1.1–numerical aperture (NA) 25× objective lens immersed in ACSF to obtain 62.5X high-magnification (166.4 μm × 166.4 μm; 512 × 512 pixels; 1.5-μm step) images for neuronal analysis. The maximum imaging depth was ◻120 μm from the pial surface. Images were acquired using low laser power (< 30 mW at the sample) and a low-pass emission filter (<700 nm).

### 2.9. Image processing and quantification

All image processing and quantification were performed with Image J software (NIH). All z-stacks of images were projected along the z-axis to recreate a two-dimensional representation of 3D structures. To quantify the microglia response to laser induced injury, we measured the number of microglia processes entering from outer area Y into inner area as a function of time. The number of red pixels in area X or Y were measured at each time point (R_x_(t) or R_y_(t), and the microglial response was defined as R(t) = [R_x_(t) − R_x_(0)](Bennett et al., 2016)/R_y_(0). To account for signal intensity differences among different experiments, we thresholded every image so that all processes had the maximum value (255), and all background was set to 0. We then counted the number of white pixels in area X over time (Rx(t)) and compared it with the first picture taken immediately after the ablation (R_x_(0)). The number of white pixels corresponds to the region covered by processes within the area X, and its increase over time provides a measure of the microglial response. To account for the variability in the number of microglia located in the outer area Y in different experiments, we calculated the microglial response relative to the number of processes in the outer area Y immediately after the ablation (R_y_(0)). The microglial response at any time point (R(t)) is therefore given by R(t) = (R_x_(t) − Rx(0))/R_y_(0).

### 2.10. Data analysis

All data are presented as mean ± s.e.m., and were analyzed by GraphPad Prism 8. Unpaired Student *t*-test and one-way ANOVA were used for group comparisons. *P* < 0.05 was considered statistically significant.

## 3. Results

### 3.1. Generation and validation of *Tmem119-tdTomato* reporter mice

Here, we took advantage of the recent identification of TMEM119 as a specific marker of human and murine microglia (Bennett et al., 2016) to generate a new tool for studying the behavior and function of microglia in health and disease. We describe the generation and the characterization of the *Tmem119-tdTomato* knock-in mouse strain where endogenous *Tmem119* expression is preserved. A CRISPR/Cas9 strategy was used to insert tdTomato, preceded by ribosome-skipping peptide porcine teschovirus-1 polyprotein (P2A), upstream of the stop codon of murine *Tmem119*. Two founder lines were established for *Tmem119-tdTomato* that were bred with wild-type mice on the C57BL/6 background to generate an on-target sequence (**Figure 1A**). The data demonstrating the specific insertion of the transgene into the *Tmem119* locus are shown (**Supplementary File 1**). Confirmation of the mouse genotypes comparing WT mice to heterozygous or homozygous lines are shown (**Figure 1B** and **Supplementary File 2**). To validate that the transgene insert does not affect endogenous Tmem119 expression, we measured *Tmem119* gene expression by quantitative PCR. As expected, we found no difference in *Tmem119* mRNA levels in brain homogenates by comparing *Tmem119*^*tdT*/+^ mice with wild-type controls (*p* = 0.9996; Student’s *t*-test; **Figure 1C**). Next, to confirm that tdTomato is co-expressed with TMEM119, we used two commercially available anti-TMEM119 antibodies (anti-intracellular and anti-extracellular domain of TMEM119) to track the TMEM119 expression in coronal brain sections from 10 months old *Tmem119*^*tdTomato*/+^ mice. Our data showed that both of the antibodies recognized the tdTomato^+^ cells, indicating that tdTomato^+^ cells readily express TMEM119.

**Figure 1.**
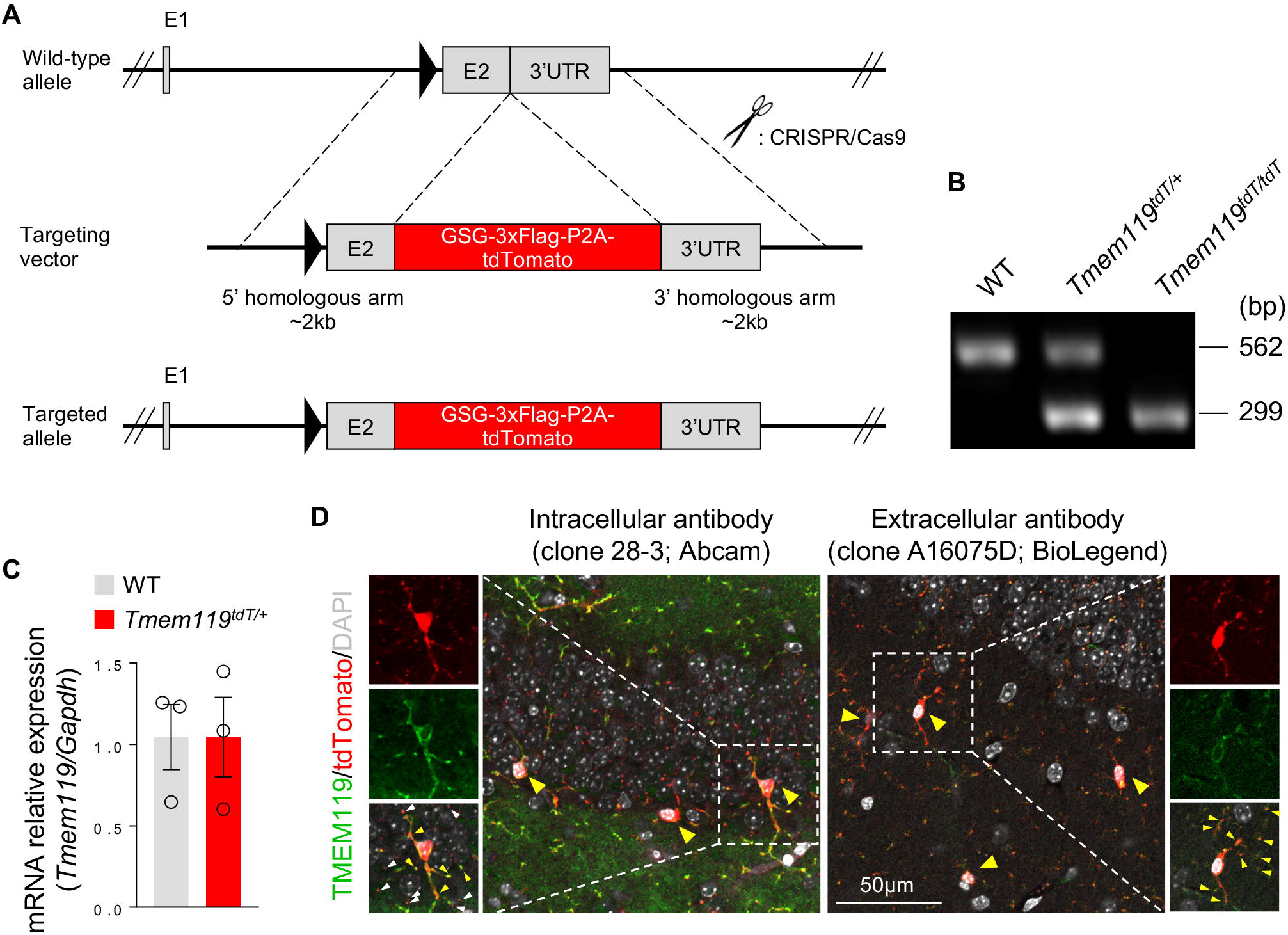
Generation and validation of *Tmem119-tdTomato* reporter mice. (**A**) Schematic of engineering strategy to create the *Tmem119-tdTomato* reporter mice. A DNA sequence of GSG-3xFlag-P2A-tdTomato was inserted between the exon 2 and 3’UTR in the *Tmem119* gene by CRISPR/Cas9 technology. E, exon; UTR, untranslated region. (**B**) A representative genotyping result showing the genotype of WT (with a 562-bp size band), *Tmem119*^tdTomato/+^ (with a 562-bp and 299-bp size band) and *Tmem119*^tdTomato/tdTomato^ (with a 299-bp band). tdT, tdTomato. (**C**) The gene expression of *Tmem119* relative to *Gapdh* in central nervous system was detected between 3-month old WT and *Tmem119*^tdTomato/+^ mice (*n* = 3 each). Data are presented in mean ± s.e.m. (**D**) TMEM119 (green) was stained in coronal brain sections from 10-month old *Tmem119*^tdTomato/+^ mice by an intracellular (left; clone 28-3, Abcam) and an extracellular (right; clone A16075D, BioLegend) antibodies. Red, tdTomato. Yellow arrows, colocalized staining. White arrows, uncolocalized staining. Scale bar, 50 μm.

### 3.2. TMEM119-tdTomato is expressed in CNS but not in peripheral tissues and blood

To examine whether tdTomato is predominately expressed in the CNS of *Tmem119-tdTomato* mice, we analyzed CNS and peripheral tissues and organs by immunofluorescence and flow cytometry. Thus, we collected brain, spinal cord, blood, heart, kidney, lung, intestine, spleen and liver from 3-month old *Tmem119*^tdTomato/+^ mice. Immunofluorescence staining showed that tdTomato positive cells were detected throughout the CNS regions, including the olfactory bulb, the prefrontal cortex, the hippocampus, the thalamus, the midbrain, the cerebellum and the spinal cord **(Figure 2A-I**). In contrast, no tdTomato expression was observed in peripheral tissues such as the heart, the kidney, the lung, the intestine, the spleen and the liver (**Figure 2J-O**). To confirm these findings, we harvested cell suspensions from these tissues as well as blood and measured the tdTomato expression by flow cytometry. Based on a gating strategy the excludes doublets and debris (**Figure 2P**), we detected 15.1% of events as tdTomato positive in the brains of *Tmem119*^tdTomato/+^ mice, but not in the brains of WT mice (**Figure 2Q**). Consistent with the histological analysis, there was no tdTomato positive cells detected in peripheral tissues (**Figure 2S-X**) or blood (**Figure 2R**).

**Figure 2.**
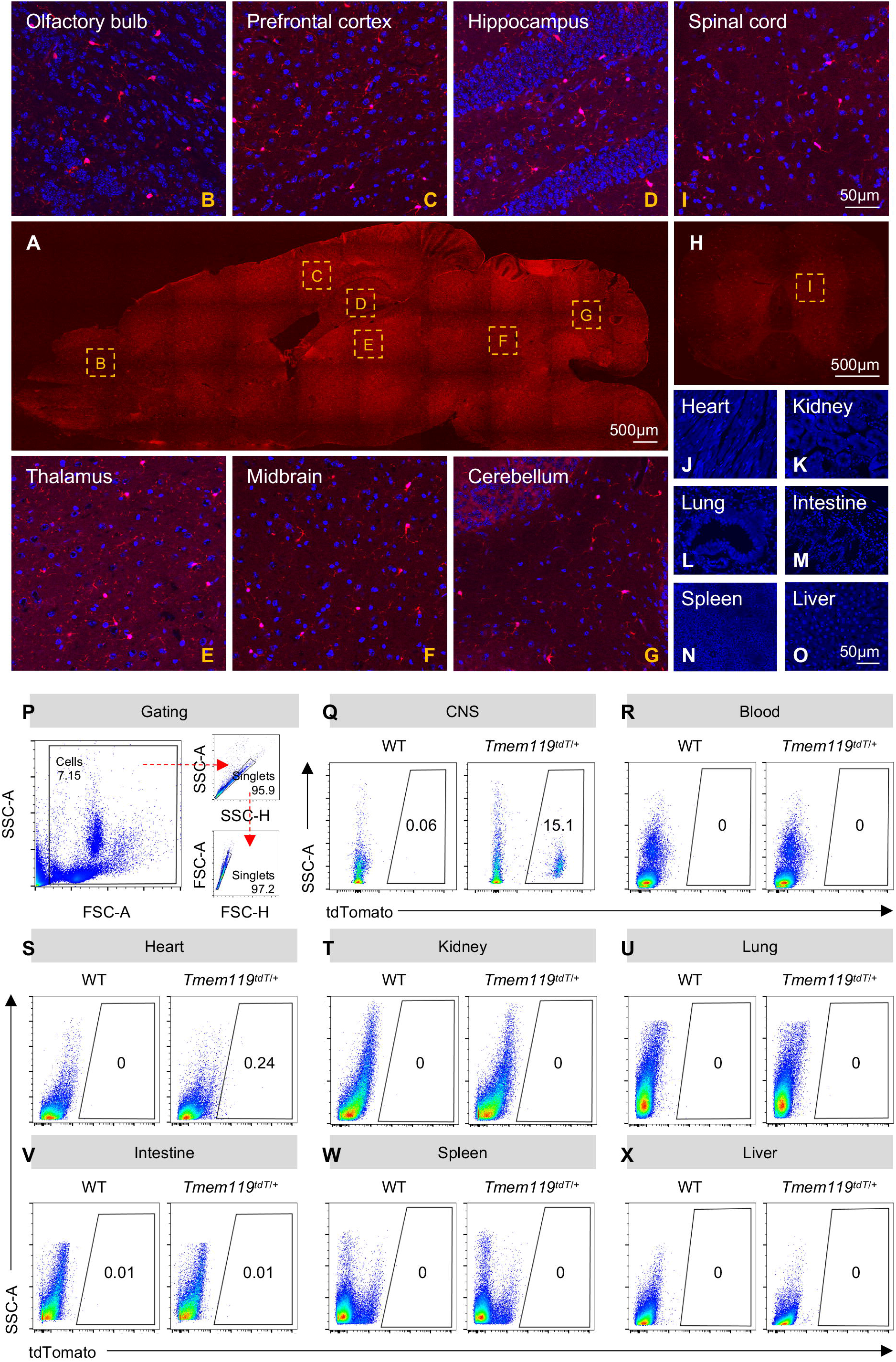
TdTomato is expressed in the adult CNS but not in the blood nor in peripheral tissues. A representative image of sagittal section in brain (**A**) and coronal section in spinal cord (H) from 3-month old *Tmem119*^tdTomato/+^ mice. Sections: 16-μm thick. Scale bar, 500 μm. TdTomato (red) were detected in cells through over different regions of central nervous system, such as olfactory bulb (**B**), prefrontal cortex (**C**), hippocampus (**D**), thalamus (**E**), midbrain (**F**), cerebellum (**G**) and spinal cord (**I**). Sections were mounted in media with DAPI (blue). Scale bar, 50 μm. TdTomato were not detected in cells in peripheral tissues such as heart (**J**), kidney (**K**), lung (**L**), intestine (**M**), spleen (**N**) and liver (**O**). Sections, 16-μm thick. Scale bar, 50 μm. TdTomato were measured in CNS, blood and peripheral tissues from 3-month old *Tmem119*^tdTomato/+^ mice by flow cytometry, and tdTomato positive cells were only detected in CNS tissues (**Q**) but not in blood (**R**) and peripheral tissues such as heart (**S**), kidney (**T**), lung (**U**), intestine (**V**), spleen (**W**) and liver (**X**), based on a gating strategy (**P**). SSC-A, side scatter (area); FSC-A, forward scatter (area); SSC-H, side scatter (hight); FSC-H, forward scatter (hight); CNS, central nervous system; WT, wild type; tdT, tdTomato.

### 3.3. TMEM119-tdTomato is constantly expressed in adult and aged CNS

We have provided evidence to prove spatial expression of TMEM119-tdTomato that is specific to the CNS. We further asked if TMEM119 expression is maintained in aged brains. To address this question, we analyzed tdTomato expression in coronal brain sections from different ages [Postnatal day (P)90, P300 and P540] of *Tmem119*^tdTomato/+^ mice by immunofluorescence staining. Our data show that TMEM119-tdTomato expression is expressed in the brain of both young adult and aged mice (**Figure 3A-C**).

**Figure 3.**
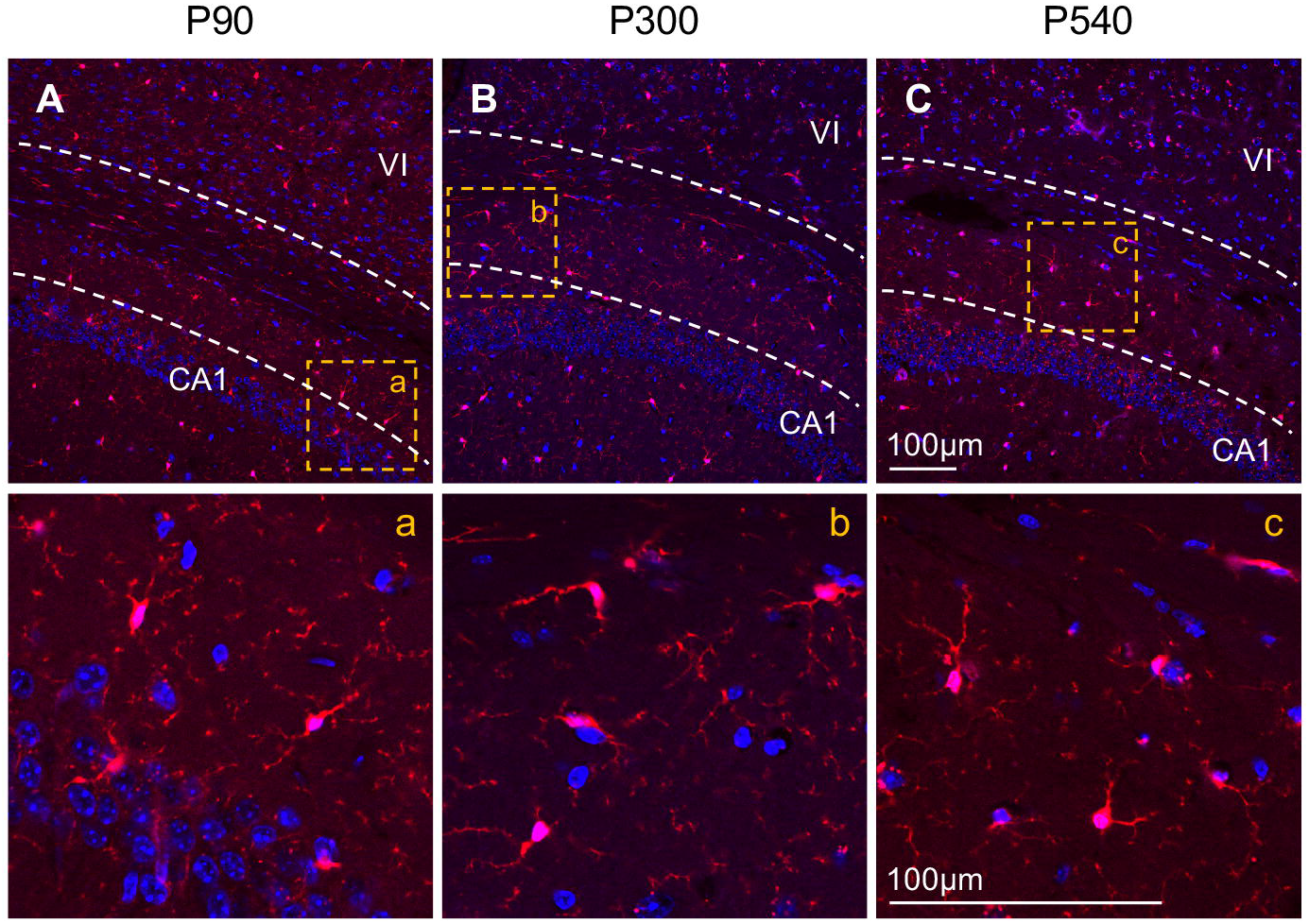
TdTomato is consistently expressed in the adult and aged CNS. TdTomato^+^ cells (red) were detectable in the coronal brain sections from postnatal 90 days (**A**), 300 days (**B**) and 540 days (**C**). **a**, **b**, and **c**, are amplified images from each staining. P, postnatal; VI, the fourth layer of neocortex; CA1, the first region of hippocampal circuit. Sections, 30-μm thick. Scale bar, 100 μm.

### 3.4. TMEM119-tdTomato is expressed in Iba1^+^ cells but not in other CNS-resident cells

Given that tdTomato is specifically expressed in microglia, we sought to examine whether TMEM119 is microglial-specific or is also detectable in other CNS-resident cells such as neurons, astrocytes and oligodendrocytes. To do this, we stained coronal brain sections of *Tmem119*^tdTomato/+^ mice with a marker of each of these three cell types in addition to a microglial marker. Thus, the following antibodies – against NeuN, GFAP, Olig2 and Iba1 - were tested in the coronal brain sections from of *Tmem119*^tdTomato/+^ mice to detect neurons, astrocytes, oligodendrocytes and microglia, respectively. Our data showed that tdTomato expression was only detected in Iba1^+^ cells but not in NeuN^+^, GFAP^+^ and Olig2^+^ cells in the brain (**Figure 4A-D**). These findings confirm that TMEM119 is exclusively expressed in microglial cells in the brain.

**Figure 4.**
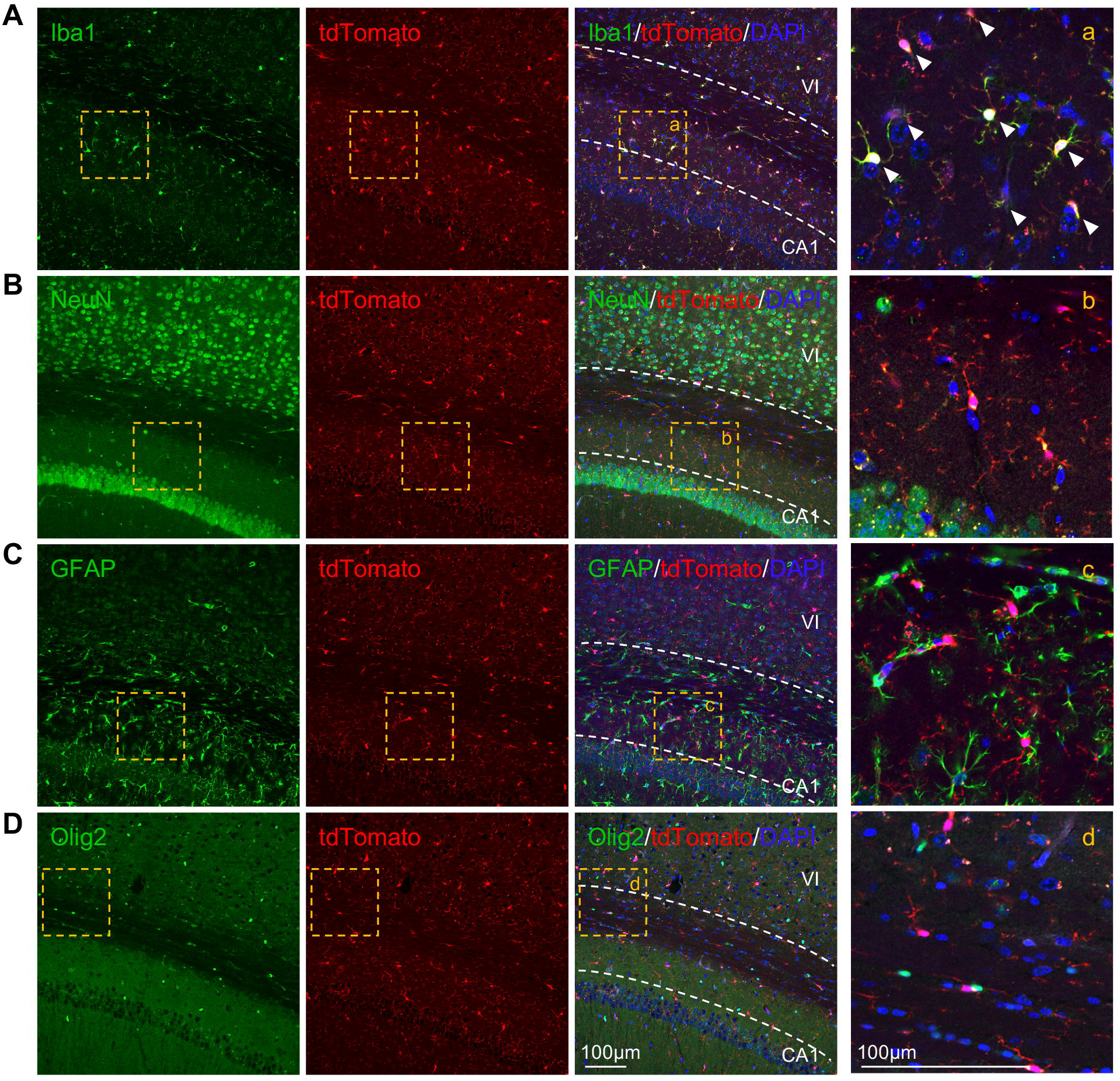
TdTomato is specifically expressed in Iba1^+^ microglia/macrophages but not in other CNS-resident cells. TdTomato^+^ cells (red) were colocalized with microglia/macrophage marker Iba1^+^ cells (**A**, green), but not neuronal marker NeuN^+^ (**B**, green), astrocyte marker GFAP^+^ (**C**, green) or oligodendrocyte marker Olig2^+^ (**D**, green) cells in the coronal brain sections of adult *Tmem119*^tdTomato/+^ mice by immunofluorescence staining. (*n* = 3 sections per mouse, *N* = 3 mice). Red, tdTomato. Blue, DAPI. **a**, **b**, **c**, and **d**, are amplified images from each staining. VI, the fourth layer of neocortex; CA1, the first region of hippocampal circuit. Arrowheads, colocalized staining. Sections, 30-μm thick. Scale bar, 100 μm.

### 3.5. TMEM119-tdTomato is specifically expressed in microglia but not in infiltrating immune cells in adult brain

To further confirm that TMEM119-tdTomato^+^ cells are indeed microglia, we used a Percoll density gradient technique to isolate microglia from the brain of adult WT and *Tmem119*^tdTomato/+^ mice. We then compared the tdTomato^+^ cells to the CD45^lo^CD11b^+^ microglia. Based on the gating strategy showed in a representative experiment in **Figure 5A-D**, we found 97.8% of live cells were CD45^+^ in WT mice (**Figure 5E**) relative to 99.4% in *Tmem119*^tdTomato/+^ mice (**Figure 5F**). Within the CD45^+^ cells, 90.6% of this population was tdTomato-positive in the *Tmem119*^tdTomato/+^ mice (**Figure 5H**), while no tdTomato was detected in WT mice (**Figure 5G**). The tdTomato^+^ cells from *Tmem119*^tdTomato/+^ mice were further analyzed, and we found that they are all CD45^lo^CD11b^+^ microglia (**Figure 5I**). In the reverse experiment, we found that 97.5% of CD45^+^ cells in WT (**Figure 5J**) and 94.6% of these cells in *Tmem119*^tdTomato/+^ mice (**Figure 5K**) are CD45^lo^CD11b^+^ microglia, and 96.6% of the CD45^lo^CD11b^+^ microglia are tdTomato positive in *Tmem119*^tdTomato/+^ mice while none are positive in WT mice (**Figure 5L**). Overall, our data demonstrate that all tdTomato^+^ cells are CD45^lo^CD11b^+^ microglia.

**Figure 5.**
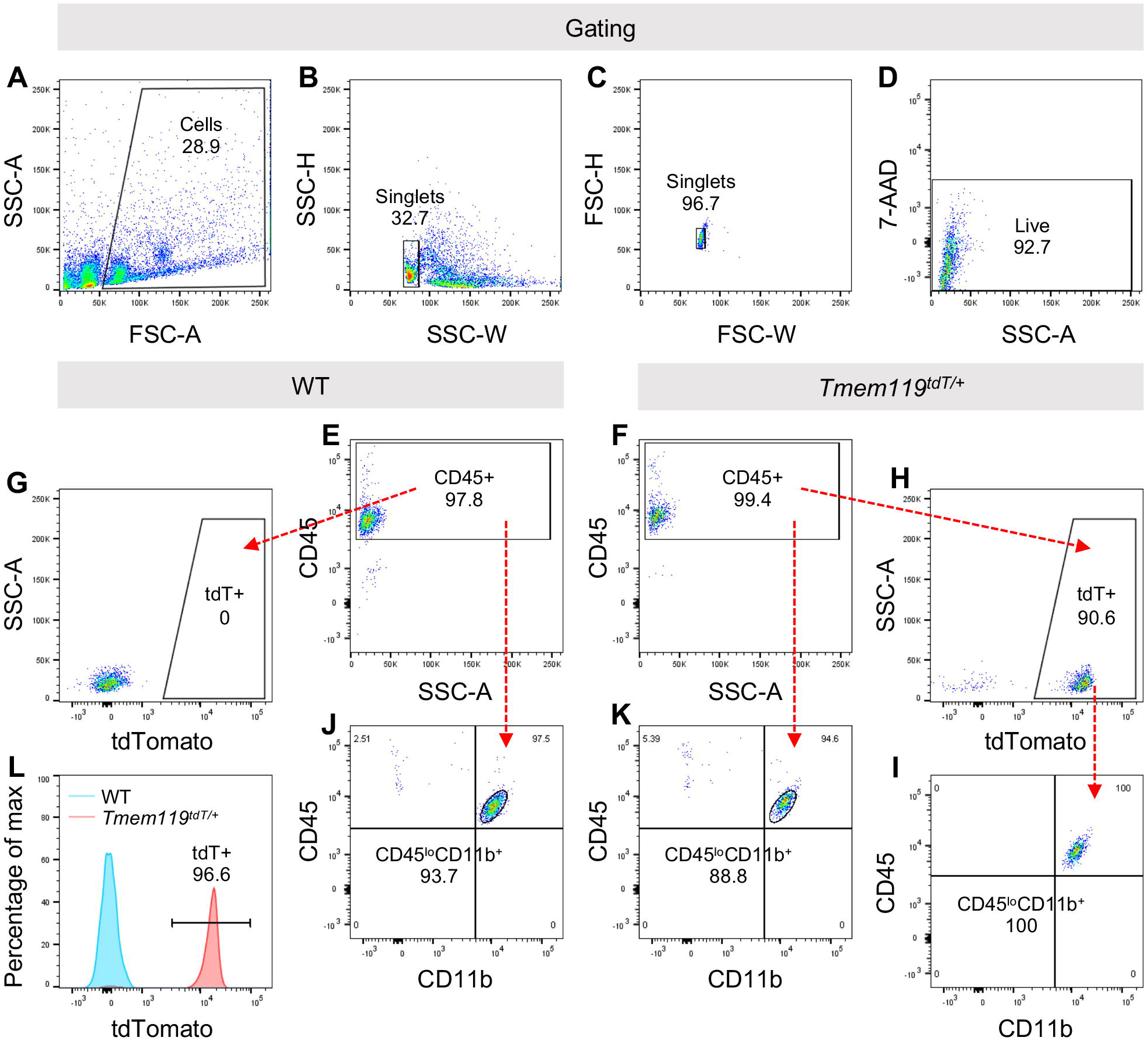
TdTomato is specifically expressed in microglia but not in infiltrating immune cells in the adult brain. (**A-D**) Flow cytometry analysis to gate out cells of interest (**A**), single cells (**B, C**) and live cells (**D**). (**E-I**) tdTomato^+^ cells in the brain tissues of WT (**G**) vs *Tmem119*^tdTomato/+^ (**H**) mice are gated based on the CD45^+^ cells in WT (**E**) or *Tmem119*^tdTomato/+^ (**F**) mice. CD45^lo^CD11b^+^ cells (**I**) were further gated out of tdTomato^+^ cells (**H**) in *Tmem119*^tdTomato/+^ mice. (**E, F, J-L**) CD45^lo^CD11b^+^ cells (**J, K**) were gated out of CD45^+^ cells in WT (**E**) or *Tmem119*^tdTomato/+^ (**F**) mice, and the tdTomato^+^ cells in WT vs *Tmem119*^tdTomato/+^ mice (**L**) were further gated based on the CD45^lo^CD11b^+^ cells (**J, K**). SSC-A, side scatter (area); SSC-H, side scatter (hight); SSC-W, side scatter (width); FSC-A, forward scatter (area); FSC-H, forward scatter (hight); FSC-W, forward scatter (width); WT, wild type; tdT, tdTomato; CD45^lo^, low expression of CD45. The number beside or within the gate indicates percentage of cells in each plot.

### 3.6. Two-photon live imaging of TMEM119-tdTomato^+^ microglia *in vivo* in a laser-induced injury model

To validate the *Tmem119-tdTomato* reporter mice as a useful tool to monitor the activity of microglia *in vivo*, we elected to use a model of laser-induced injury to monitor microglia response in the intact living brain. Using transcranial two-photon microscopy, we were able to image the behavior of tdTomato-expressing parenchymal microglia through the open skull window of anesthetized *Tmem119*^tdTomato/+^ mice. Thus we performed a small laser ablation, ~30 μm in diameter, inside the primary sensory cortex (**Figure 6A**). Time-lapse images (up to150 minutes) show that microglia formed a spherical containment around the laser lesion site. Within minutes after laser injury, the number of microglia entering from outer area (Y) into inner area (X) is measured as a function of time showing rapid response of microglia to the injury site (**Figure 6B, C**). During this period, the same cells also retracted those processes that previously lay in directions opposite to the site of injury. Most of the cellular content of each of the immediate neighbors was directed towards the damaged site within the first 1-3 h (**Figure 6D**, **Supplementary Video 1**). Length changes of four processes marked in **Figure 6D** were quantified as function of time (**Figure 6E**). These results indicate that the *Tmem119-tdTomato* mouse strain is a viable model to monitor and analyze the activity and the formation of microglia processes in neuroinflammatory disease models.

**Figure 6.**
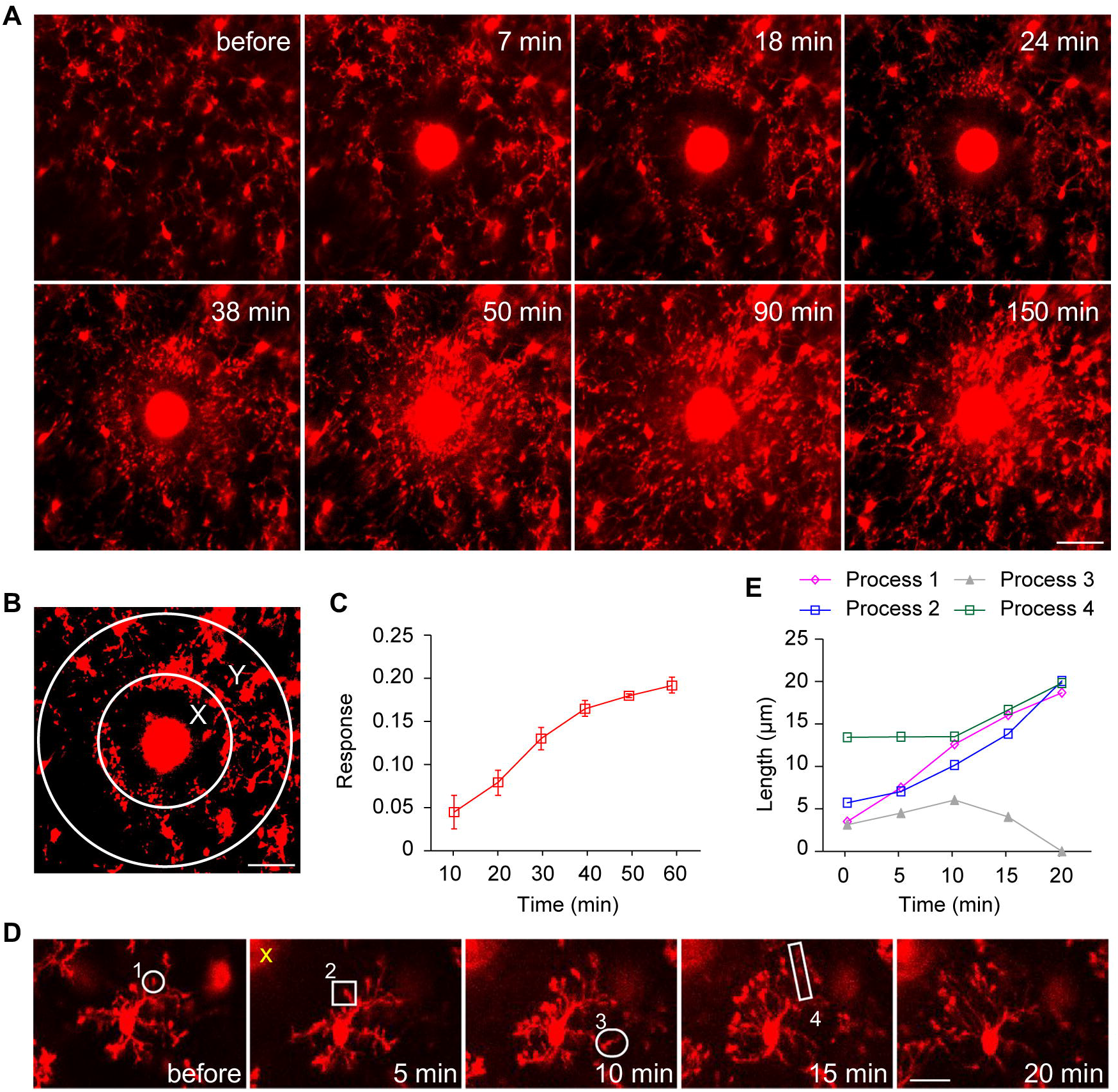
Two-photon live imaging of tdTomato^+^ microglia *in vivo* after laser-mediated injury. (**A**) After a localized ablation inside the primary sensory cortex, neighboring Tmem119-tdTonmato^+^ microglia respond quickly with extended processes and bulbous termini. Time-lapse images show that microglia formed a spherical containment around the laser lesion site. Scale bar, 30 μm. (**B**) To quantify the laser injury-induced microglial response, the number of microglia processes entering from outer area Y into inner area X is measured as a function of time. The number of red pixels in area X or Y were measured at each time point (R_x_(t) or R_y_(t), and the microglial response was defined as R(t) = [R_x_(t) − R_x_(0)]/R_y_(0). Scale bar, 30 μm. (**C**) Microglia response over 1 hour after laser injury. (**D**) The same cell extends its processes towards the laser lesion site (marked by yellow x), and retracts the ones in the opposite direction to the injured site. Scale bar, 10 μm. (**E**) Length changes of four processes marked in **D** as a function of time. Scale bar, 10 μm.

## 4. Discussion

Microglia are becoming the center of attention in the field of neurodegeneration and several clinical trials are currently designed to manipulate microglial function in the context of neurodegenerative diseases (Fu et al., 2019; Mullard, 2018; Subramaniam and Federoff, 2017). However, cultured microglia quickly lose their unique microglia signature once out of the CNS, making the ability to study these cells difficult. Here, we present a model which allows us to study microglial behavior *in vivo*, even in the presence of other innate immune cells. Given the lack of markers that discriminate between microglia and infiltrated myeloid cells that are shown to invade the CNS in neurodegenerative diseases, developing new tools to detect and isolate microglia is becoming of high interest to the research community. In the present study, we report a microglia-specific mouse strain that express the recently identified microglia-specific marker Tmem119 (Bennett et al., 2016; Satoh et al., 2016). A knock-in strategy was used to generate the *Tmem119-tdTomato* where *tdTomato* is placed between exon 2 and the 3’UTR of *Tmem119* gene thus allowing the detection of TMEM119-positive cells without affecting TMEM119 endogenous expression as shown by comparing Tmem119 expression in the brains of *Tmem119-tdTomato* and wild-type mice. The choice of tdTomato as a reporter protein is because it provides a very bright red fluorescence (~3 fold higher than EGFP)(Shaner et al., 2005) that is ideal for live imaging and potentially for generating double reporter mice with GFP mice for studying microglia in relation to other GFP-labeled cell types.

Using immunofluorescence and flow cytometry analysis we provide evidence that tdTomato selectively labels microglial cells in the CNS but not peripheral myeloid cells. Satoh *et al*. showed that TMEM119 could discriminate brain-residing microglial cells from circulating macrophages, as this protein is expressed exclusively on ramified and amoeboid microglial cells in the brain (Satoh et al., 2016). Additionally, we validate the specificity of two commercially available anti-TMEM119 antibodies available at Abcam and Biolegend that stain the intracellular and extracellular domains of TMEM119, respectively. In adult Tmem119-tdTomato reporter mice, we found that TMEM119 is expressed throughout the brain and the spinal cord. Strikingly, we did not detect the tdTomato signal in peripheral lymphoid organs and the blood. Bennett and colleagues generated monoclonal antibodies to the intracellular and extracellular domains of TMEM119 that enable the immunolabeling of microglia in histological sections as well as isolation of microglia by FACS (Bennett et al., 2016). The study by Bennett *et al.* showed that Tmem119 immunoreactivity is developmentally regulated as Tmem119 expression did not appear until postnatal day 3 and 6 (Bennett et al., 2016). These findings are in agreement with the present study where we did not detect tdTomato expression in the brain of newborn mice (data not shown).

During the past few years and with the availability of instrumental gene profiling techniques that can process very small numbers of cells, mainly from human autopsies and biopsies, led to a series of reports generating transcriptomics profiling of human and murine microglia at the bulk and the single-cell levels (Butovsky et al., 2014; Chiu et al., 2013; Hickman et al., 2013; Marta Olah, 2019; Masuda et al., 2019; Olah et al., 2018; Zhang et al., 2014). Several microglia clusters have been identified with specific gene signatures that are associated with homeostatic and active microglia. In our recent study examining the gene signature of the aged human brain using fresh brain autopsies from aged subjects, we found that *Tmem119* gene expression is found in microglia purified from aged human brain (Olah et al., 2018). This is consistent with our present data in the *Tmem119-tdTomato* mice where TMEM119 expression is readily present in aged microglia (1.6 years old mice) at the protein level. More recently, given the heterogeneous nature of the human brain, we have utilized a single cell RNA-seq approach to analyze microglia heterogeneity across several neurological disorders including Alzheimer’s disease, Parkinson’s disease and epilepsy. Computational analysis identified nine distinct microglia subsets that express different levels of *Tmem119* (Marta Olah, 2019). In our *Tmem119-tdTomato* mice, although TMEM119 expression was detected throughout the healthy adult and aged brains, it remains possible that TMEM119 expression will be regulated in a disease state where microglia are subject to neuroinflammatory stimuli. Another recent study reporting single-cell RNA-seq analyses of microglia in mice and humans showed that Tmem119 is among the most highly differentially regulated genes during development as it is highly expressed in postnatal microglia. The authors found that Tmem119 is expressed in homeostatic microglia clusters in healthy brains whereas microglia clusters that are associated with multiple sclerosis and are characterized as activated clusters express lower levels of Tmem119 (Masuda et al., 2019). Single-cell RNA-seq approaches are an instrumental approach to define heterogeneous cell subsets. Crossing the Tmem119-tdTomato mice with mouse models of neurodegenerative diseases such as Alzheimer’s or Parkinson’s disease will provide valuable information about the different subsets of TMEM119-positive microglia in a disease state.

In addition to the goals discussed above that lead to the generation of microglia-specific reporter mice based on the TMEM119 approach, we were interested in having in hand a reporter model where we can image microglia *in vivo* in live animals. Our two-photon imaging data demonstrate that the *Tmem119-tdTomato* mice are an excellent model for live observation of microglia to analyze their response to any type of stimulation or stress as reported using laser-induced injury. Indeed, our two-photon analysis reported the number of tdTomato-positive microglia processes entering the site of injury, processes formation and their retraction vis-à-vis the site of injury. Our findings suggest that the *Tmem119-tdTomato* mouse strain is an excellent model for imaging and analyzing microglia activity alone or in relation to other CNS-resident or infiltrated immune cells in disease models such as multiple sclerosis, Parkinson’s disease and Alzheimer’s disease.

In conclusion, our study provides a microglia-specific reporter mouse model based on the Tmem119 microglia marker. This *Tmem119-tdTomato* mouse strain will be a critical tool for microglia-focused research in health and disease. Additionally, crossing this mouse strain with other reporter mice expressing other fluorescent proteins specific for CNS-resident cells and peripheral immune cells will provide unique mouse models to study the interaction between microglia and other CNS resident or infiltrated cells.

## Acknowledgments

We thank Pfizer for the generous gift of the *Tmem119-tdTomato* reporter mice. Research reported in this publication was supported by a grant from the Thompson Family Foundation (W.E.).

## Supplementary data

**Supplementary file 1: Detailed workflow for the design and the generation of the *Tmem119-tdTomato* knock-in mice.** Five sections are provided to highlight the workflow of the mouse generation starting from the targeting strategy using CRISPR/Cas9 technology, the Cas9/sgRNA plasmid contruction and UCA assay, targeting vector construction & Southern blot strategy design, sgRNA preparation, the zygote microinjection & founder genotyping.

**Supplementary file 2: Genotyping guide of the *Tmem119-tdTomato* knock-in mice.** This report includes primer design strategy, primer sequences and predicted product sizes, PCR reaction system & conditions, gel images, and determination of genotype from the gel image results.

**Supplementary video: Time-lapse video of TMEM119-tdTomato^+^ microglia response to laser-induced injury.** After a localized ablation inside the primary sensory cortex, neighboring tdTomato^+^ microglia respond quickly with extended processes and bulbous termini.

