## Supplementary material for "A Novel *Tmem119-tdTomato* Reporter Mouse Model for Studying Microglia in the Central Nervous System": Supplemetary File 1

### Slide 1
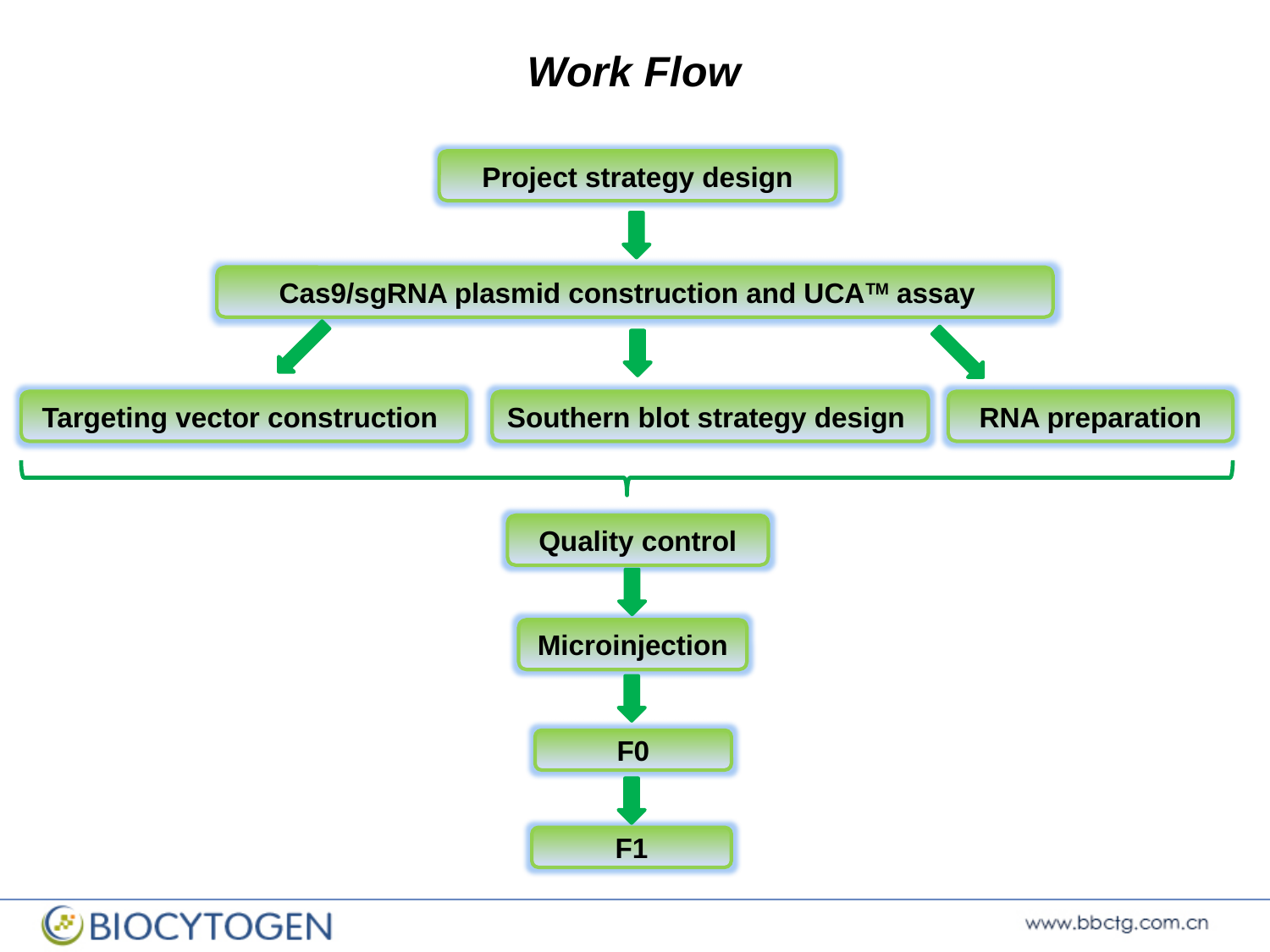

Work Flow
Project strategy design
Cas9/sgRNA plasmid construction and UCA^TM^ assay
Targeting vector construction
Southern blot strategy design
RNA preparation
Quality control
Microinjection
F0
F1

### Slide 2
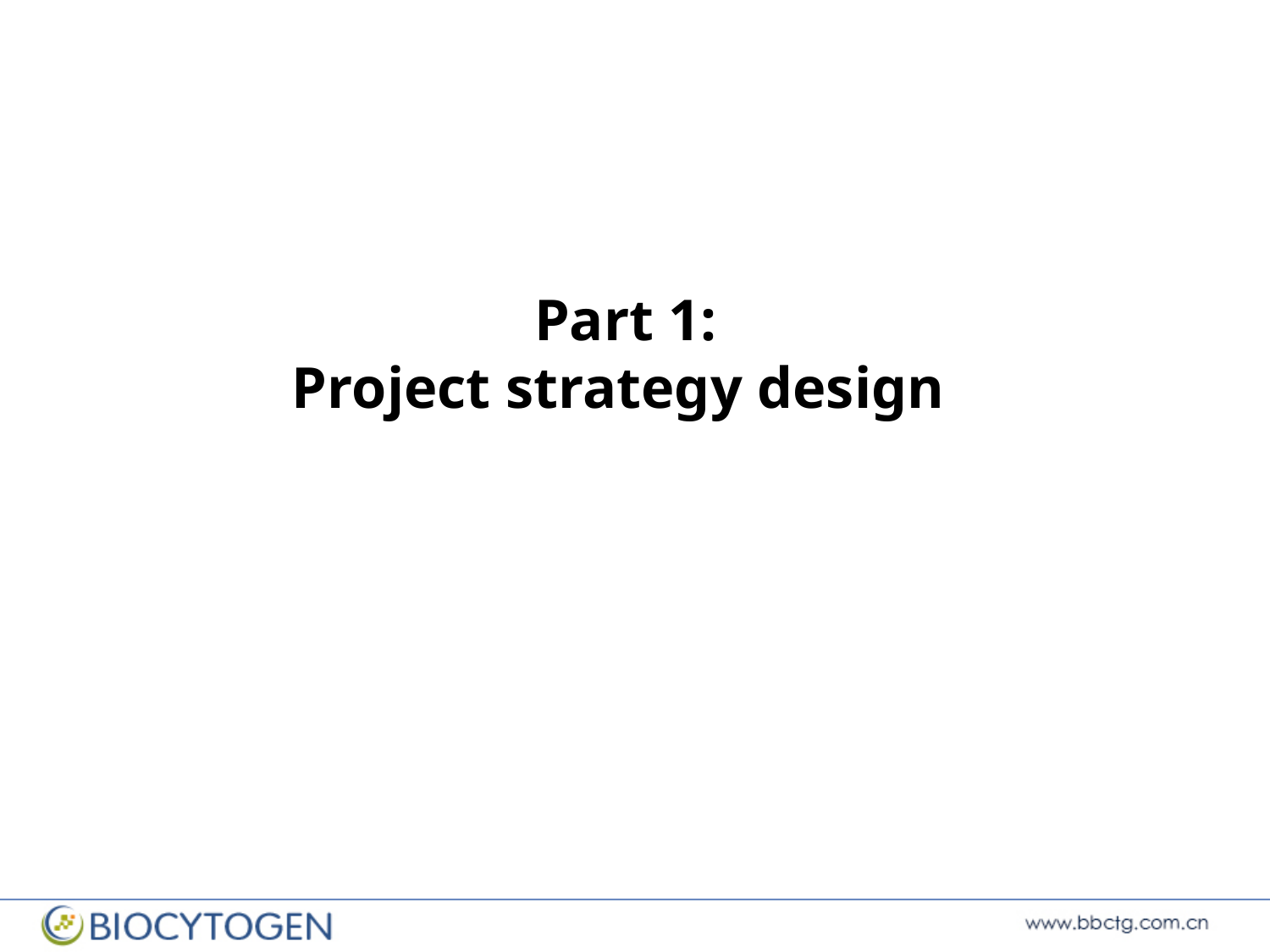

Part 1:
Project strategy design

### Slide 3
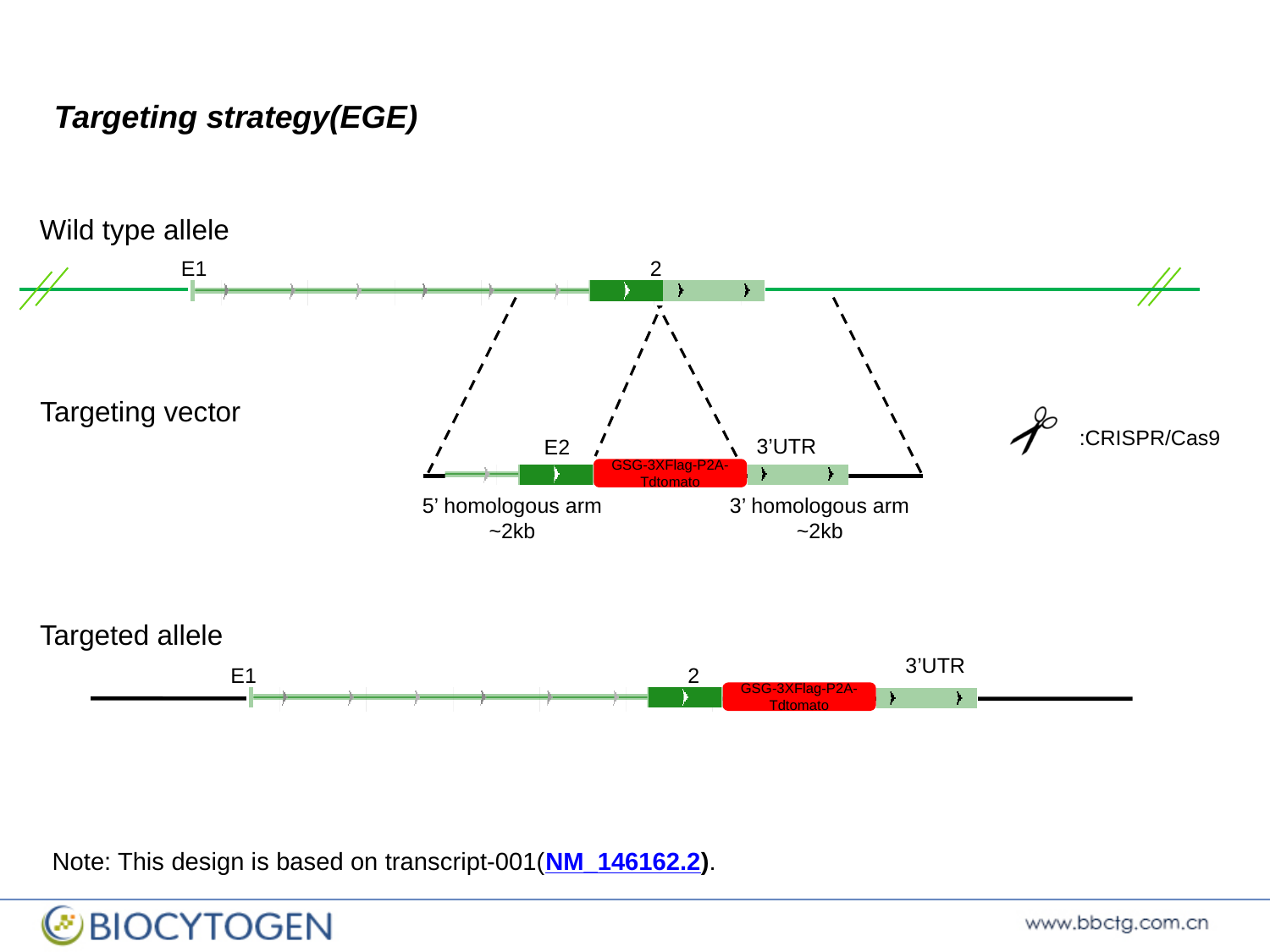

Targeting strategy(EGE)
Wild type allele
E1 2
Targeting vector
:CRISPR/Cas9
3’UTR
 E2
GSG-3XFlag-P2A-Tdtomato
5’ homologous arm
~2kb
3’ homologous arm
~2kb
Targeted allele
3’UTR
E1 2
GSG-3XFlag-P2A-Tdtomato
Note: This design is based on transcript-001(NM_146162.2).

### Slide 4
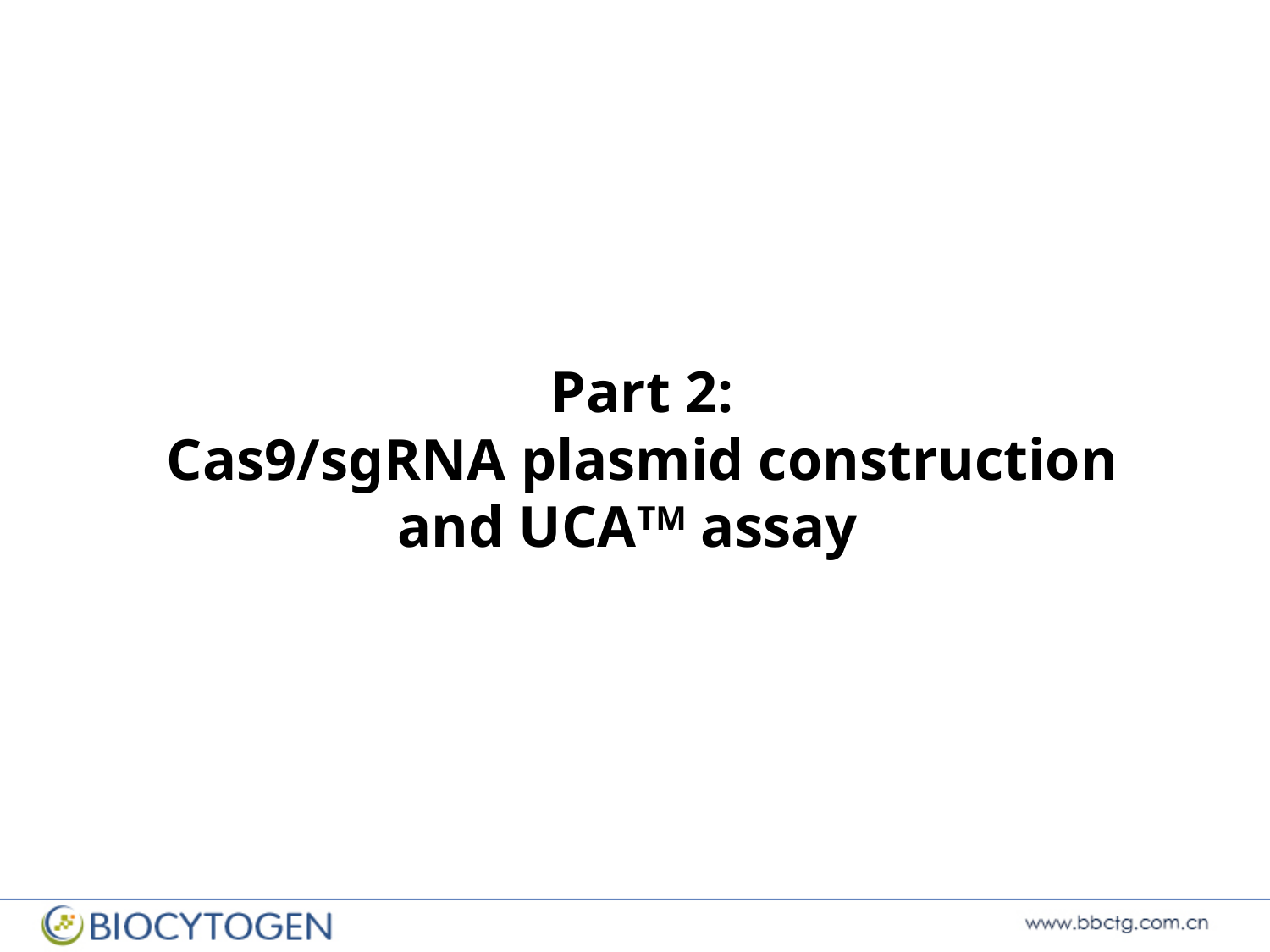

Part 2:
Cas9/sgRNA plasmid construction and UCA^TM^ assay

### Slide 5
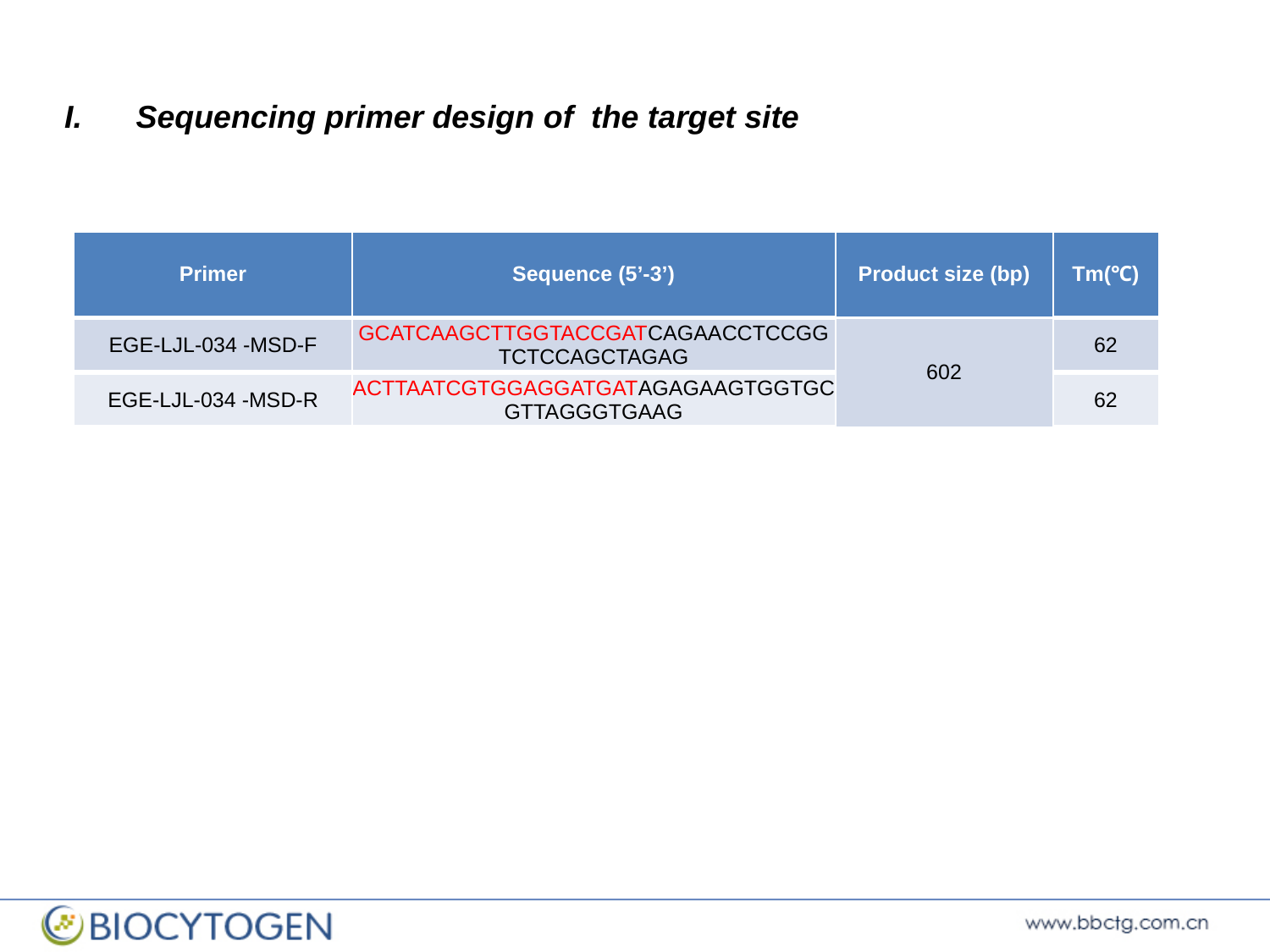

Sequencing primer design of the target site
| Primer | Sequence (5’-3’) | Product size (bp) | Tm(℃) |
| --- | --- | --- | --- |
| EGE-LJL-034 -MSD-F | GCATCAAGCTTGGTACCGATCAGAACCTCCGGTCTCCAGCTAGAG | 602 | 62 |
| EGE-LJL-034 -MSD-R | ACTTAATCGTGGAGGATGATAGAGAAGTGGTGCGTTAGGGTGAAG | | 62 |

### Slide 6
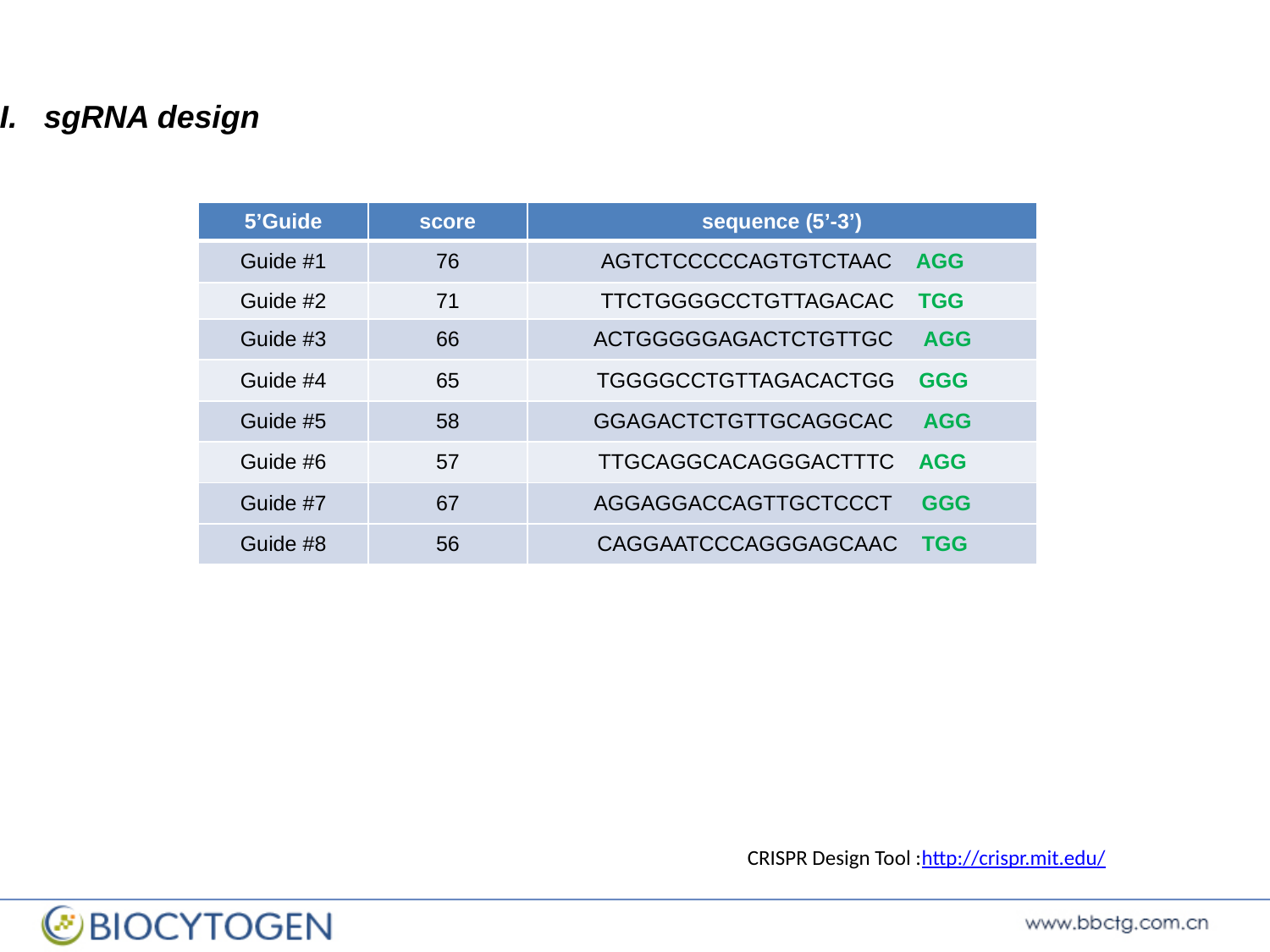

II. sgRNA design
| 5’Guide | score | sequence (5’-3’) |
| --- | --- | --- |
| Guide #1 | 76 | AGTCTCCCCCAGTGTCTAAC AGG |
| Guide #2 | 71 | TTCTGGGGCCTGTTAGACAC TGG |
| Guide #3 | 66 | ACTGGGGGAGACTCTGTTGC AGG |
| Guide #4 | 65 | TGGGGCCTGTTAGACACTGG GGG |
| Guide #5 | 58 | GGAGACTCTGTTGCAGGCAC AGG |
| Guide #6 | 57 | TTGCAGGCACAGGGACTTTC AGG |
| Guide #7 | 67 | AGGAGGACCAGTTGCTCCCT GGG |
| Guide #8 | 56 | CAGGAATCCCAGGGAGCAAC TGG |
CRISPR Design Tool :http://crispr.mit.edu/

### Slide 7
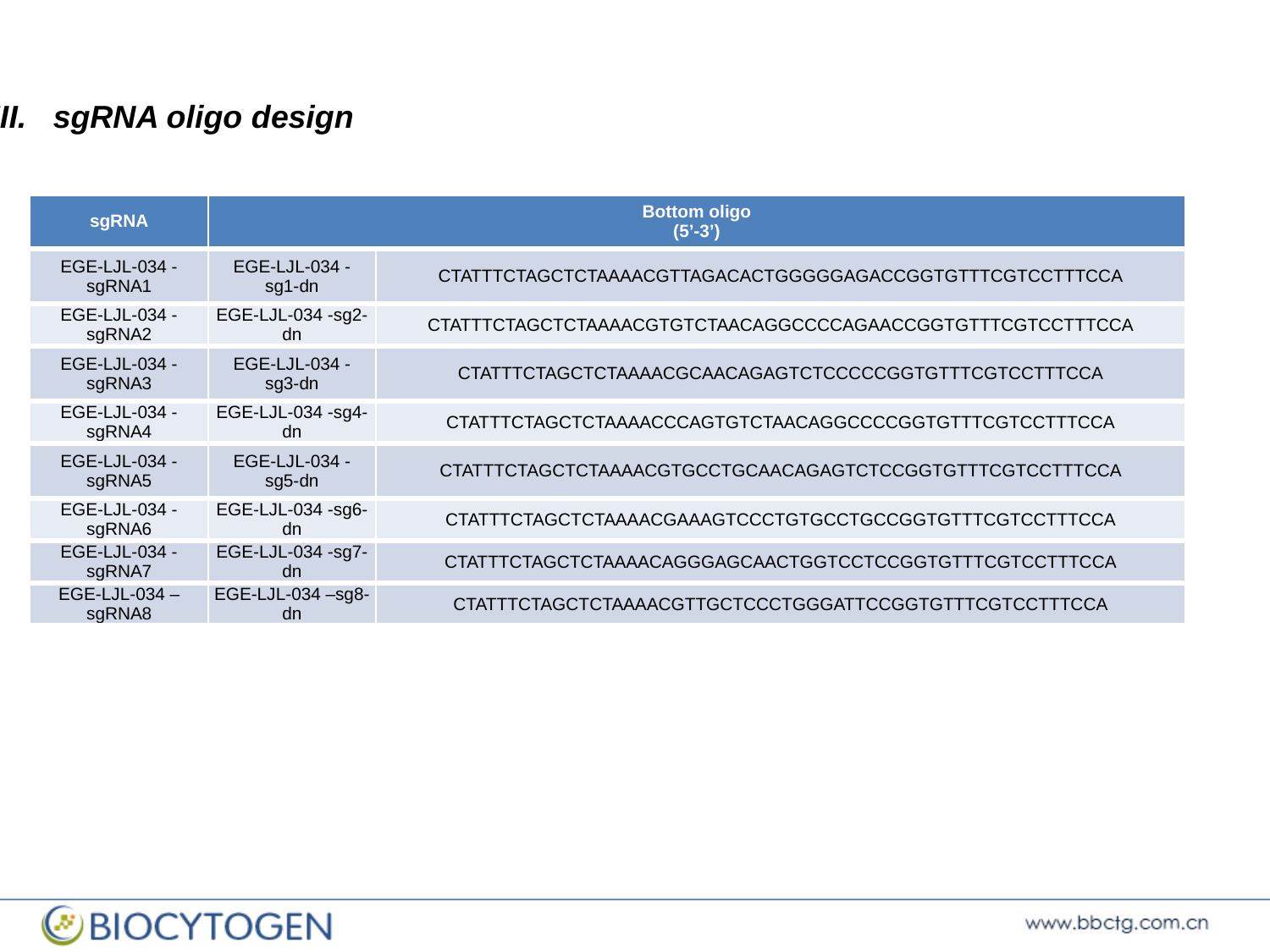

III. sgRNA oligo design
| sgRNA | Bottom oligo (5’-3’) | |
| --- | --- | --- |
| EGE-LJL-034 -sgRNA1 | EGE-LJL-034 -sg1-dn | CTATTTCTAGCTCTAAAACGTTAGACACTGGGGGAGACCGGTGTTTCGTCCTTTCCA |
| EGE-LJL-034 -sgRNA2 | EGE-LJL-034 -sg2-dn | CTATTTCTAGCTCTAAAACGTGTCTAACAGGCCCCAGAACCGGTGTTTCGTCCTTTCCA |
| EGE-LJL-034 -sgRNA3 | EGE-LJL-034 -sg3-dn | CTATTTCTAGCTCTAAAACGCAACAGAGTCTCCCCCGGTGTTTCGTCCTTTCCA |
| EGE-LJL-034 -sgRNA4 | EGE-LJL-034 -sg4-dn | CTATTTCTAGCTCTAAAACCCAGTGTCTAACAGGCCCCGGTGTTTCGTCCTTTCCA |
| EGE-LJL-034 -sgRNA5 | EGE-LJL-034 -sg5-dn | CTATTTCTAGCTCTAAAACGTGCCTGCAACAGAGTCTCCGGTGTTTCGTCCTTTCCA |
| EGE-LJL-034 -sgRNA6 | EGE-LJL-034 -sg6-dn | CTATTTCTAGCTCTAAAACGAAAGTCCCTGTGCCTGCCGGTGTTTCGTCCTTTCCA |
| EGE-LJL-034 -sgRNA7 | EGE-LJL-034 -sg7-dn | CTATTTCTAGCTCTAAAACAGGGAGCAACTGGTCCTCCGGTGTTTCGTCCTTTCCA |
| EGE-LJL-034 –sgRNA8 | EGE-LJL-034 –sg8-dn | CTATTTCTAGCTCTAAAACGTTGCTCCCTGGGATTCCGGTGTTTCGTCCTTTCCA |

### Slide 8
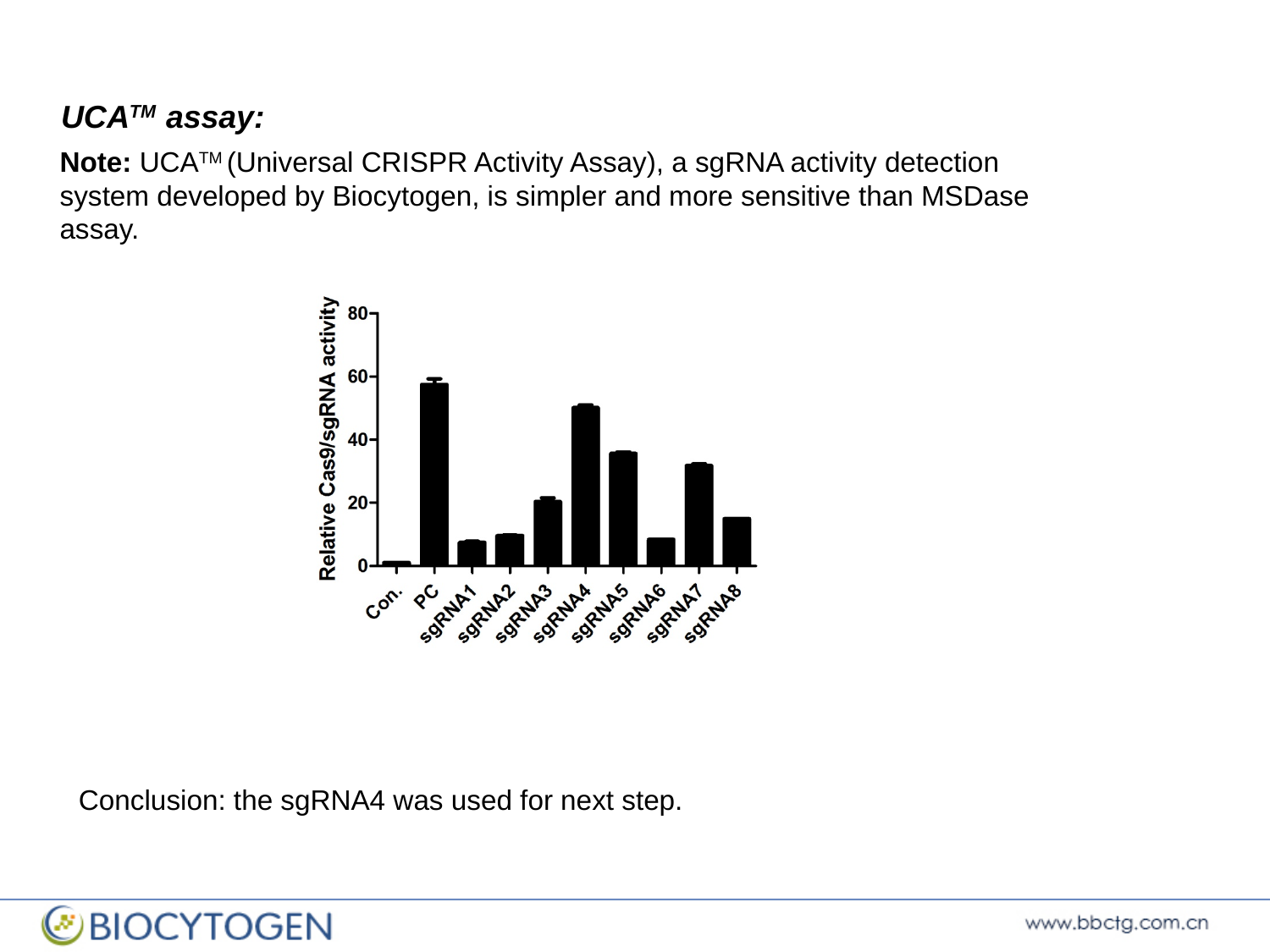

UCA^TM^ assay:
Note: UCA^TM^ (Universal CRISPR Activity Assay), a sgRNA activity detection system developed by Biocytogen, is simpler and more sensitive than MSDase assay.
Conclusion: the sgRNA4 was used for next step.

### Slide 9
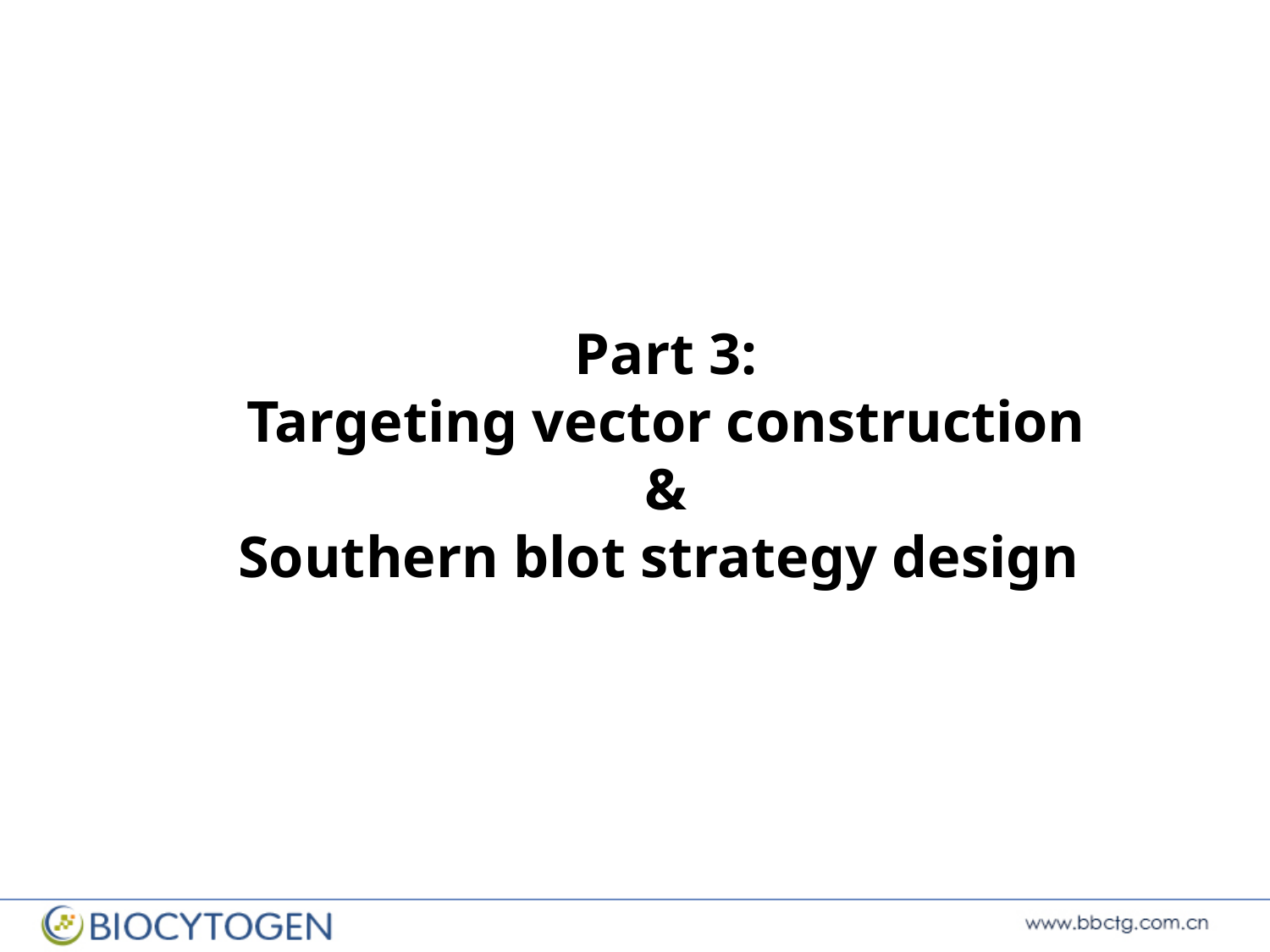

Part 3:
Targeting vector construction
&
Southern blot strategy design

### Slide 10
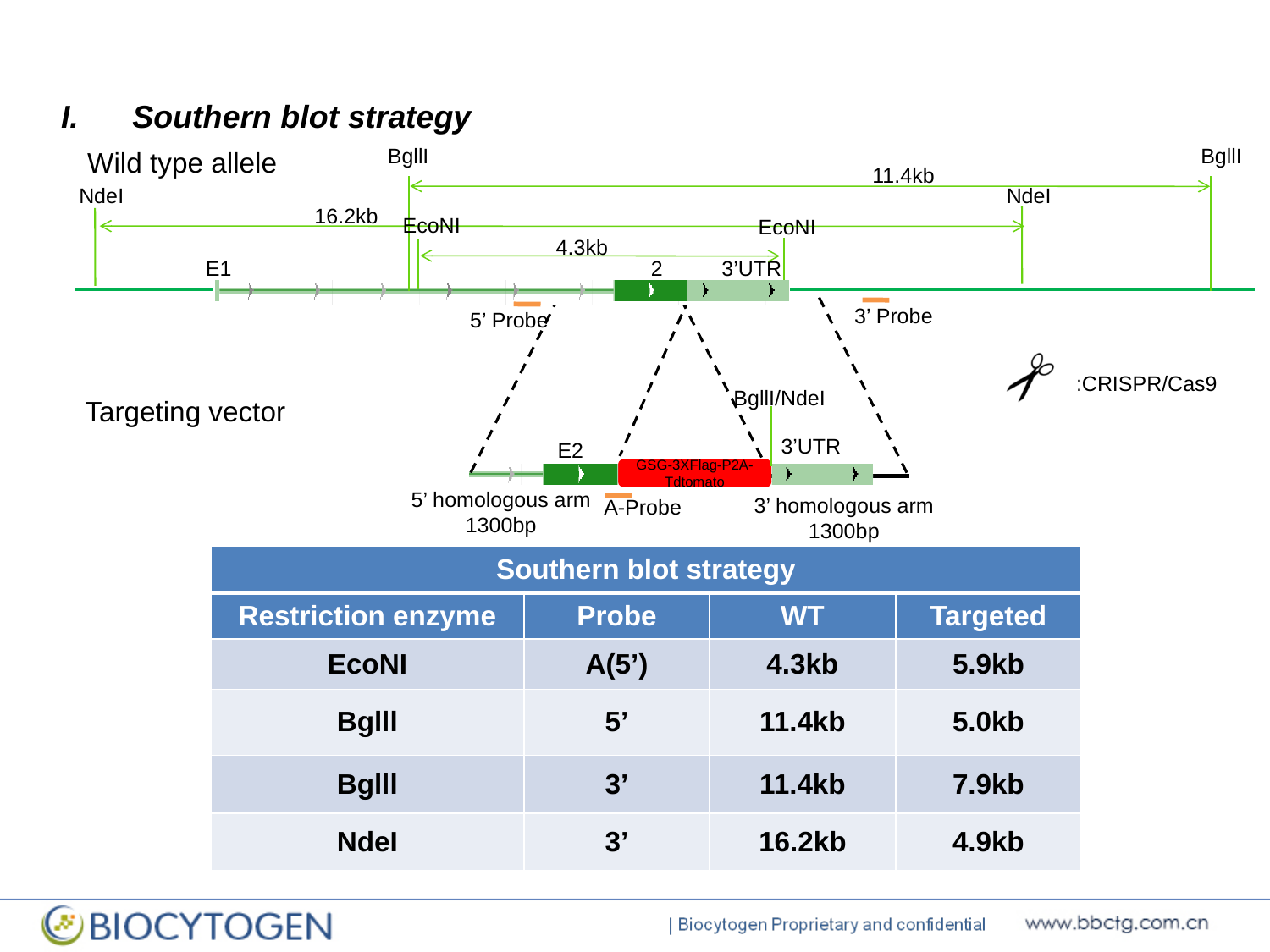

Southern blot strategy
BgllI
BgllI
Wild type allele
11.4kb
NdeI
NdeI
16.2kb
EcoNI
EcoNI
4.3kb
E1 2 3’UTR
3’ Probe
5’ Probe
:CRISPR/Cas9
BgllI/NdeI
 Targeting vector
 E2
3’UTR
GSG-3XFlag-P2A-Tdtomato
5’ homologous arm
1300bp
3’ homologous arm
1300bp
A-Probe
| Southern blot strategy | | | |
| --- | --- | --- | --- |
| Restriction enzyme | Probe | WT | Targeted |
| EcoNI | A(5’) | 4.3kb | 5.9kb |
| Bglll | 5’ | 11.4kb | 5.0kb |
| Bglll | 3’ | 11.4kb | 7.9kb |
| NdeI | 3’ | 16.2kb | 4.9kb |

### Slide 11
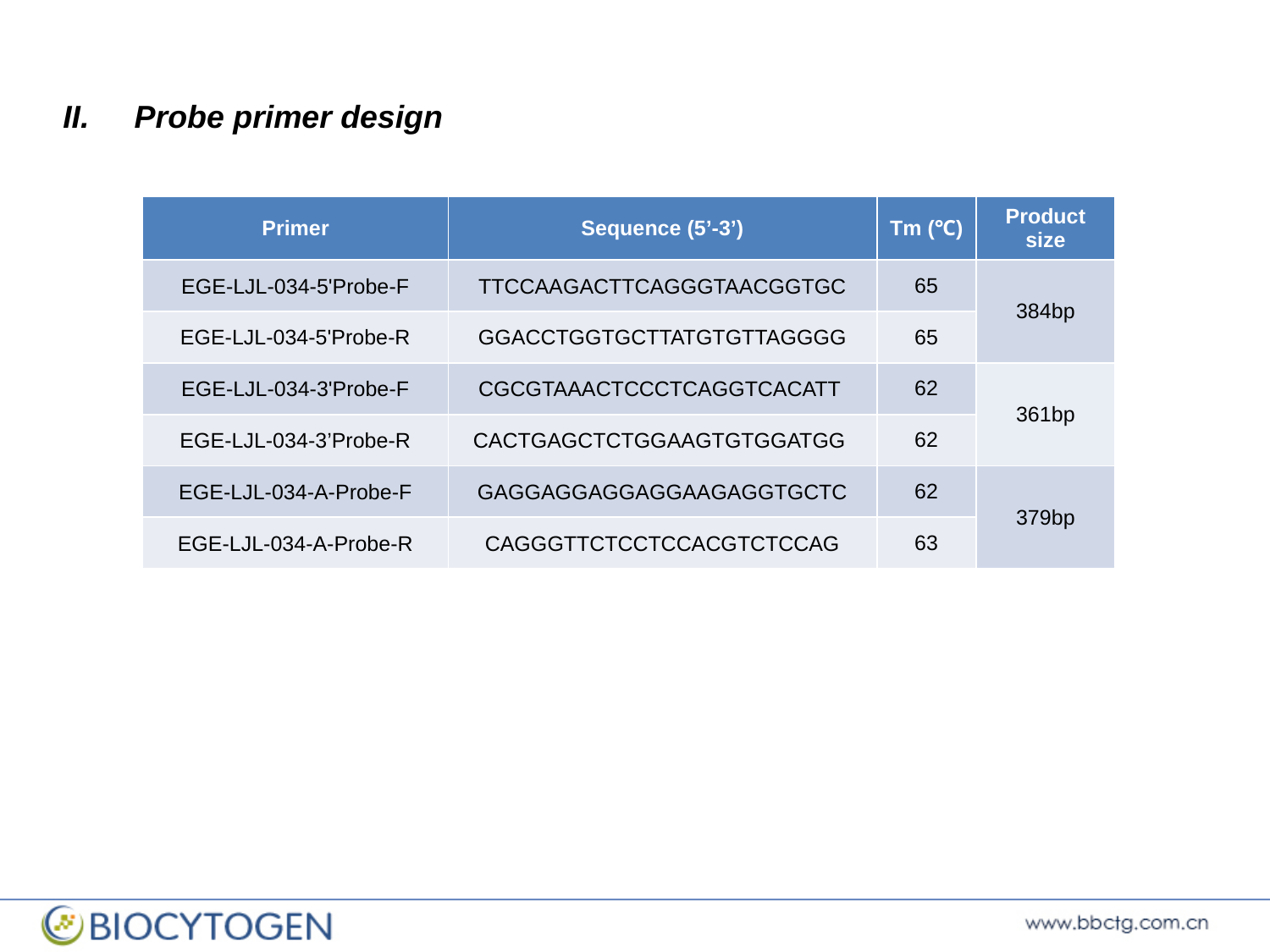

Probe primer design
| Primer | Sequence (5’-3’) | Tm (℃) | Product size |
| --- | --- | --- | --- |
| EGE-LJL-034-5'Probe-F | TTCCAAGACTTCAGGGTAACGGTGC | 65 | 384bp |
| EGE-LJL-034-5'Probe-R | GGACCTGGTGCTTATGTGTTAGGGG | 65 | |
| EGE-LJL-034-3'Probe-F | CGCGTAAACTCCCTCAGGTCACATT | 62 | 361bp |
| EGE-LJL-034-3’Probe-R | CACTGAGCTCTGGAAGTGTGGATGG | 62 | |
| EGE-LJL-034-A-Probe-F | GAGGAGGAGGAGGAAGAGGTGCTC | 62 | 379bp |
| EGE-LJL-034-A-Probe-R | CAGGGTTCTCCTCCACGTCTCCAG | 63 | |

### Slide 12
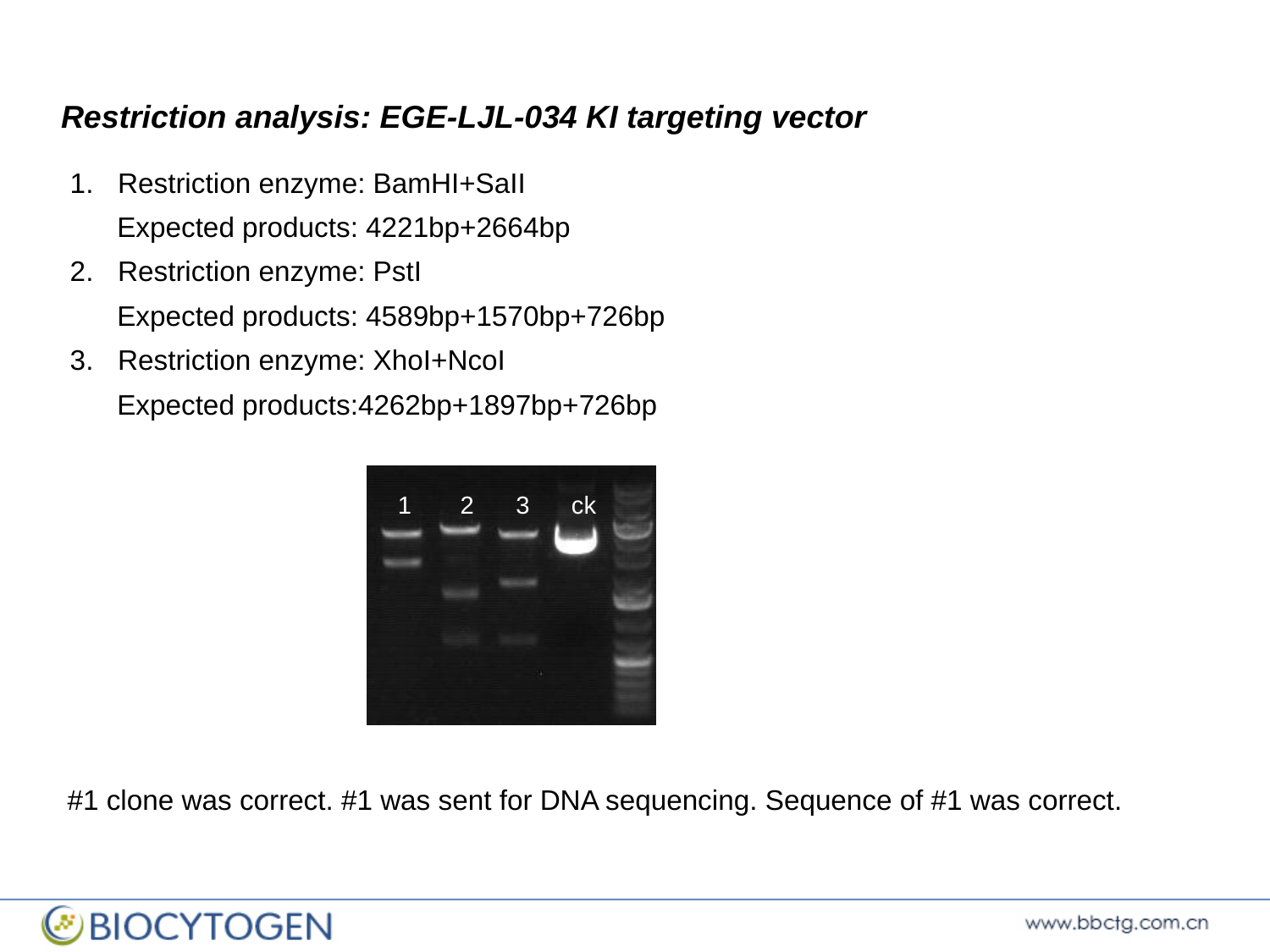

Restriction analysis: EGE-LJL-034 KI targeting vector
Restriction enzyme: BamHI+SaII
 Expected products: 4221bp+2664bp
Restriction enzyme: PstI
 Expected products: 4589bp+1570bp+726bp
Restriction enzyme: XhoI+NcoI
 Expected products:4262bp+1897bp+726bp
1 2 3 ck
#1 clone was correct. #1 was sent for DNA sequencing. Sequence of #1 was correct.

### Slide 13
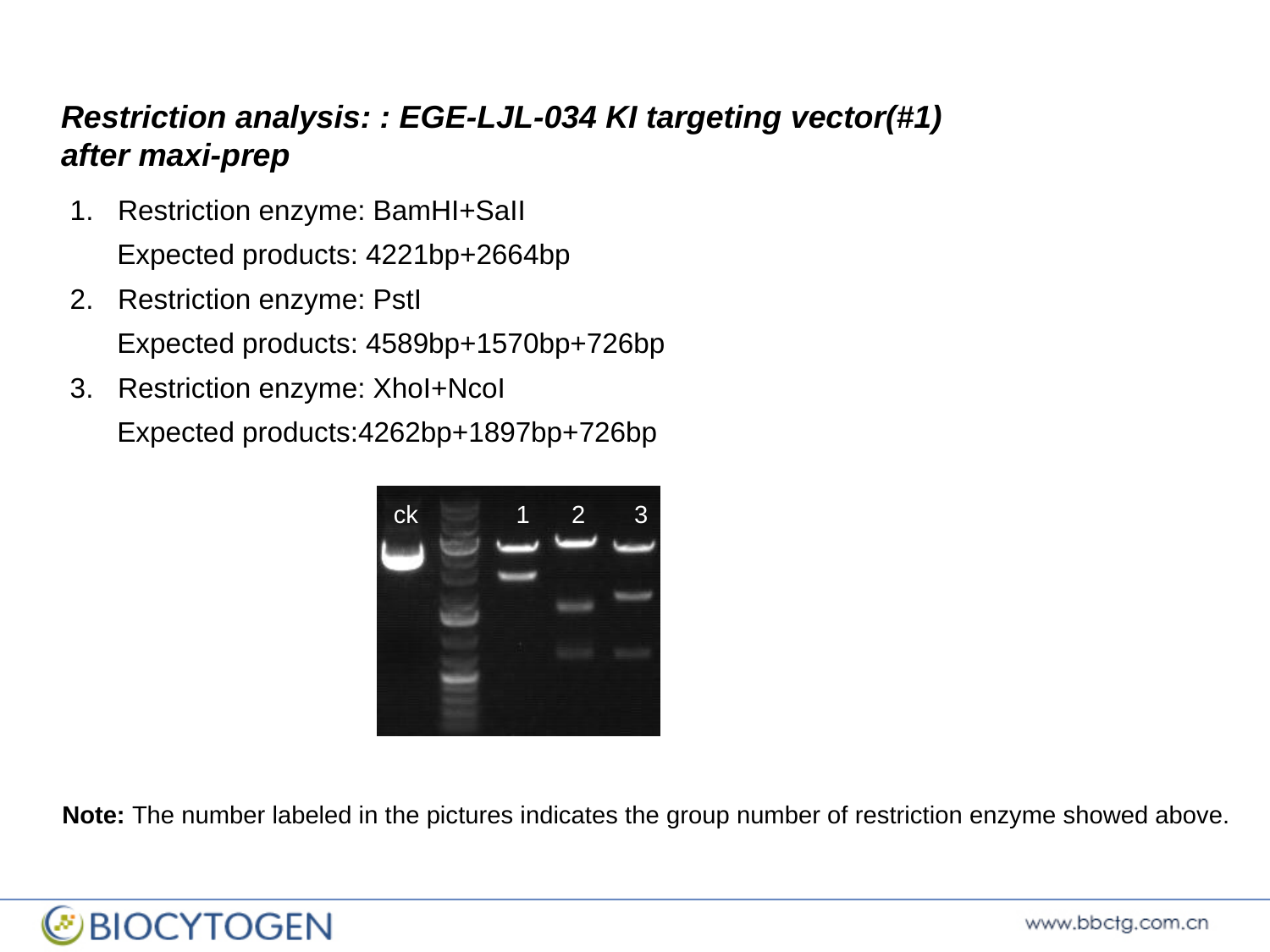

Restriction analysis: : EGE-LJL-034 KI targeting vector(#1) after maxi-prep
Restriction enzyme: BamHI+SaII
 Expected products: 4221bp+2664bp
Restriction enzyme: PstI
 Expected products: 4589bp+1570bp+726bp
Restriction enzyme: XhoI+NcoI
 Expected products:4262bp+1897bp+726bp
 ck 1 2 3
Note: The number labeled in the pictures indicates the group number of restriction enzyme showed above.

### Slide 14
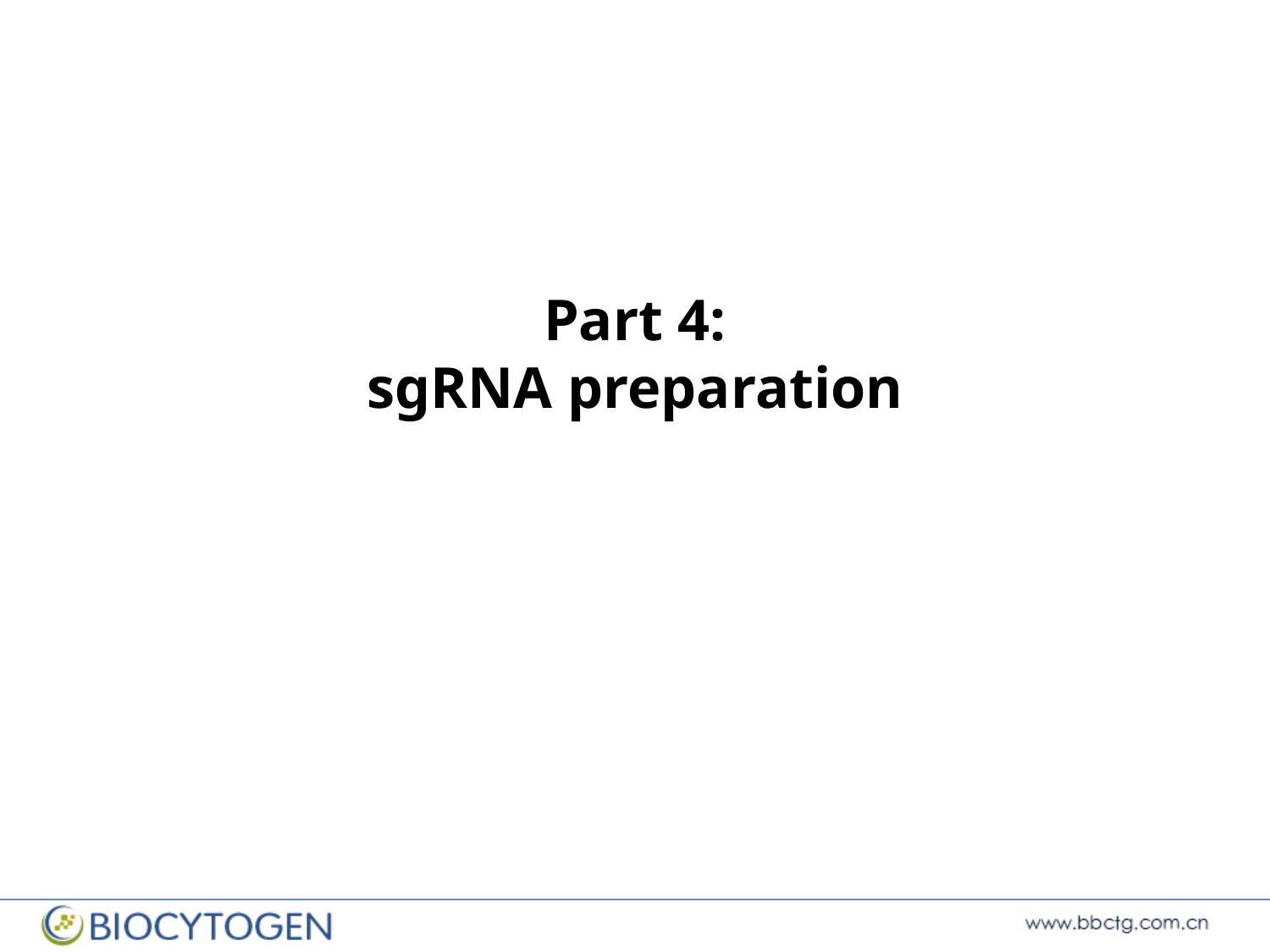

Part 4:
sgRNA preparation

### Slide 15
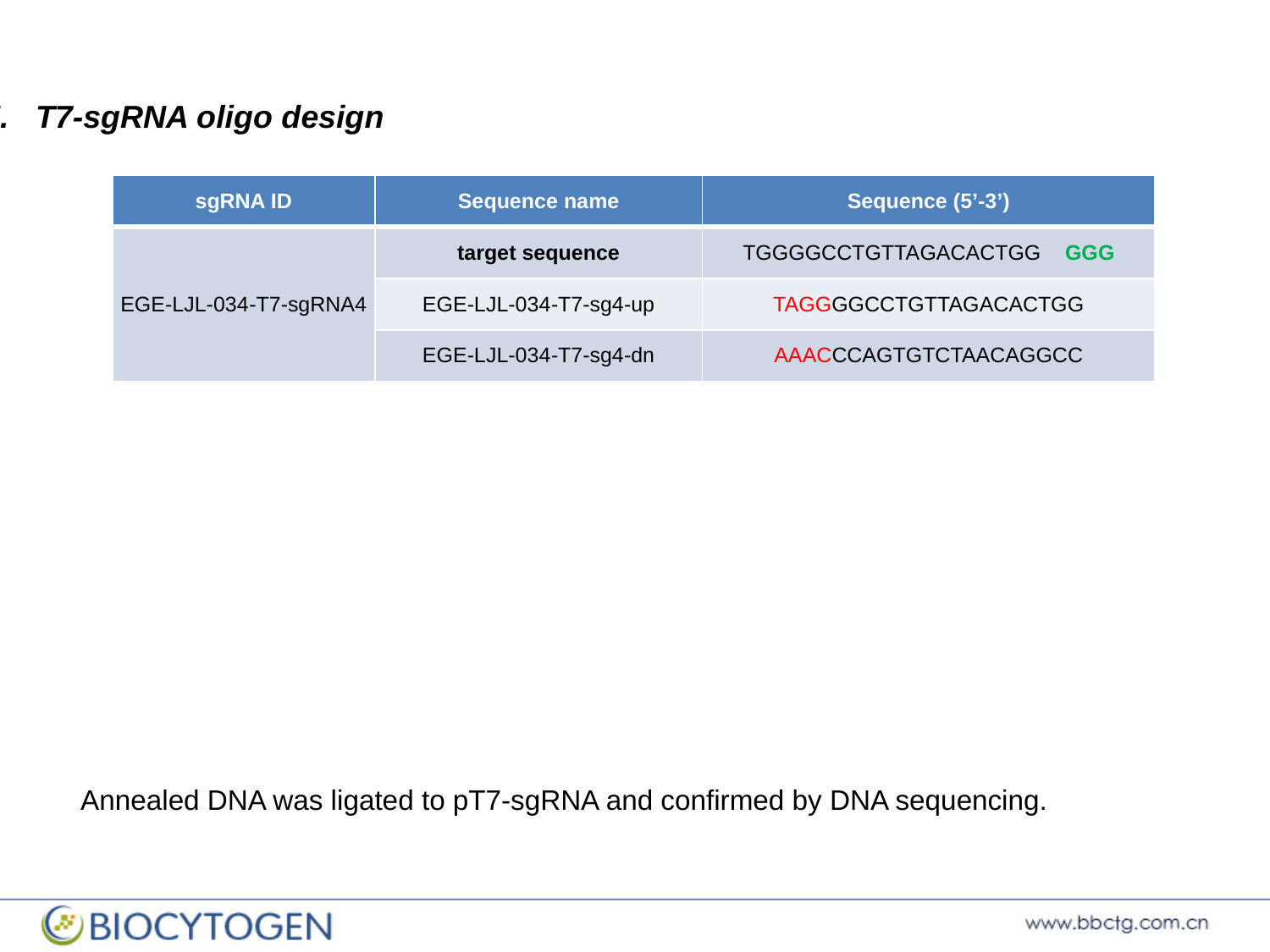

I. T7-sgRNA oligo design
| sgRNA ID | Sequence name | Sequence (5’-3’) |
| --- | --- | --- |
| EGE-LJL-034-T7-sgRNA4 | target sequence | TGGGGCCTGTTAGACACTGG GGG |
| | EGE-LJL-034-T7-sg4-up | TAGGGGCCTGTTAGACACTGG |
| | EGE-LJL-034-T7-sg4-dn | AAACCCAGTGTCTAACAGGCC |
Annealed DNA was ligated to pT7-sgRNA and confirmed by DNA sequencing.

### Slide 16
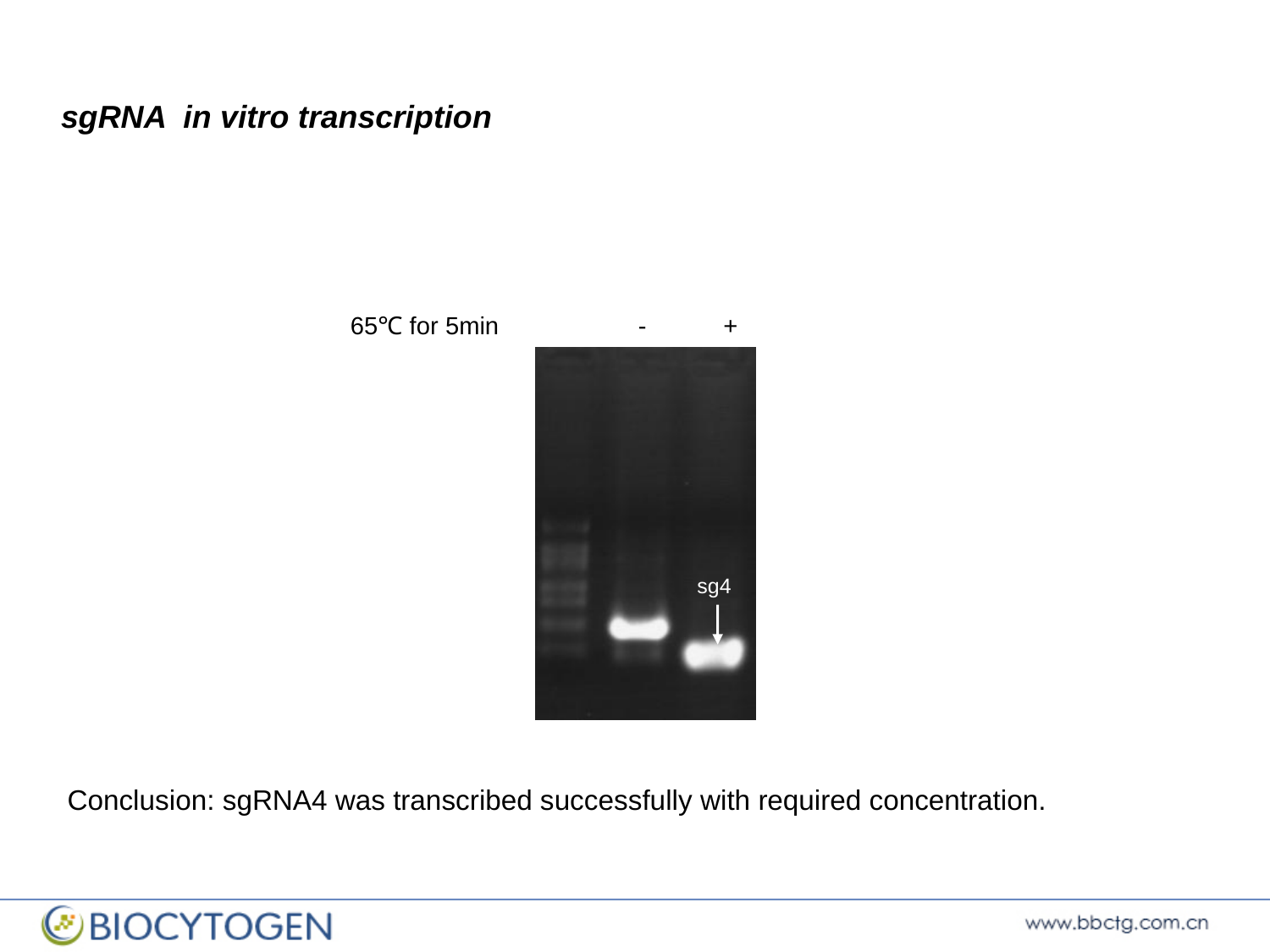

sgRNA in vitro transcription
 65℃ for 5min - +
sg4
Conclusion: sgRNA4 was transcribed successfully with required concentration.

### Slide 17
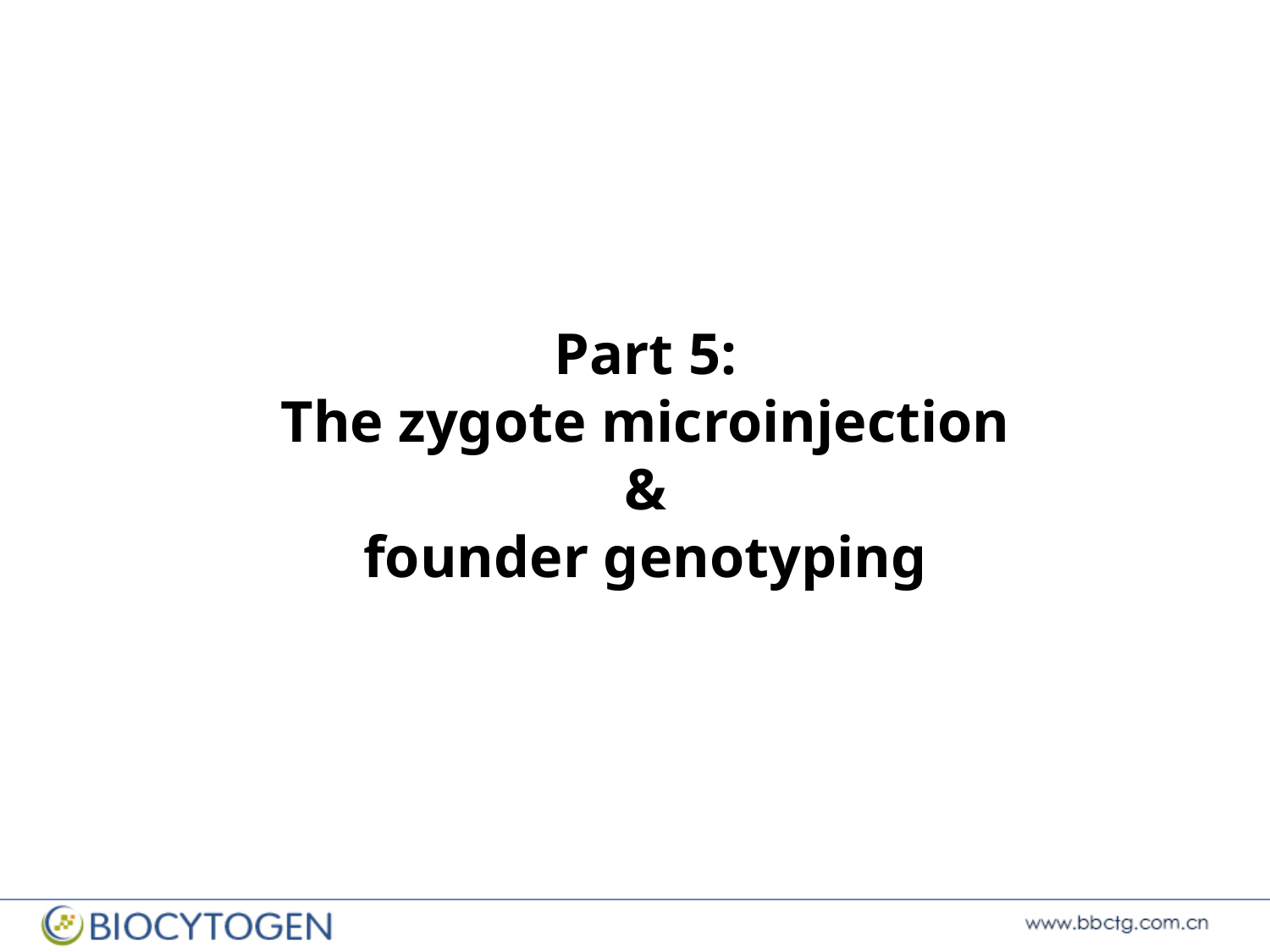

Part 5:
The zygote microinjection
&
founder genotyping

### Slide 18
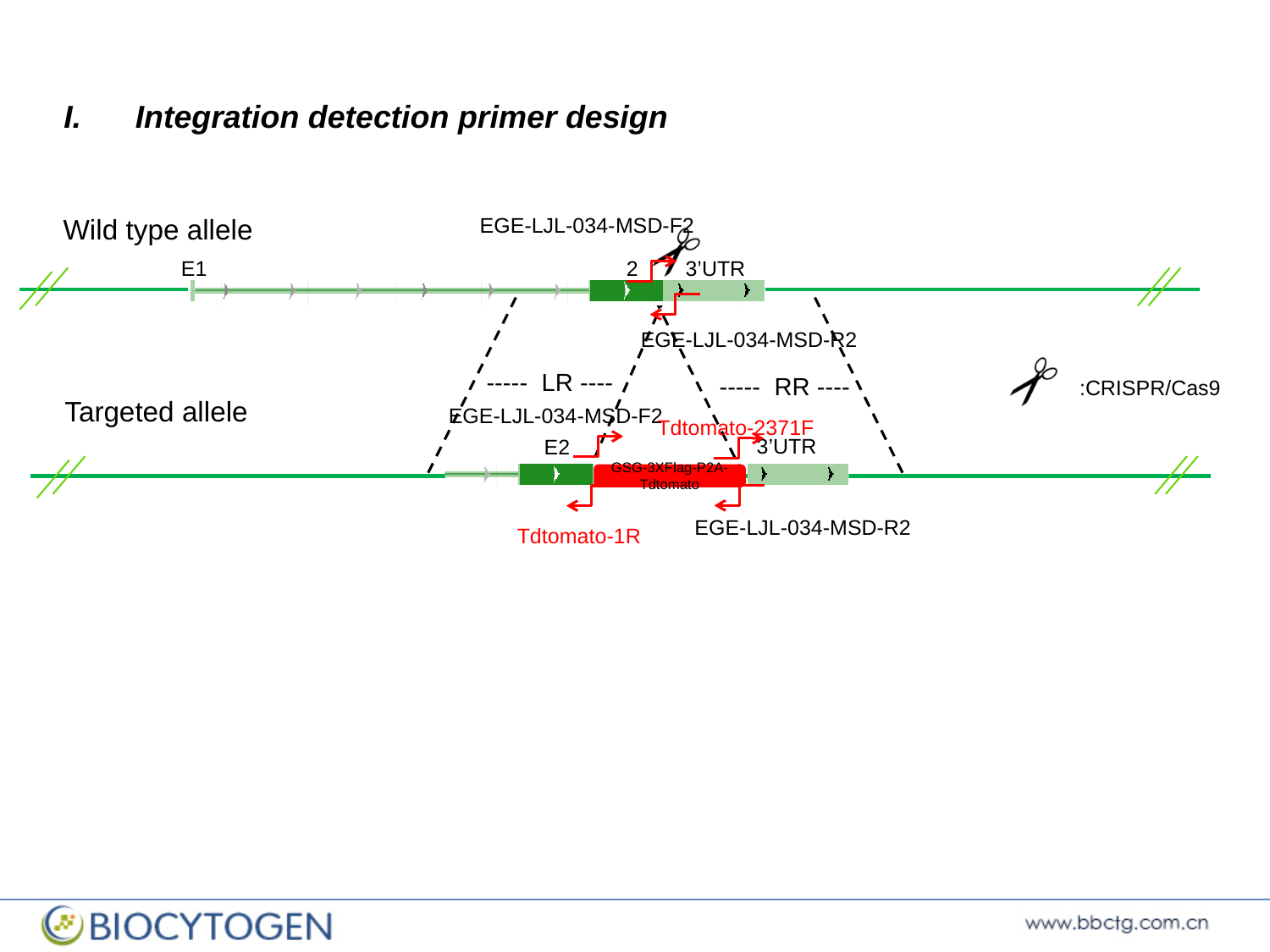

Integration detection primer design
Wild type allele
EGE-LJL-034-MSD-F2
E1 2 3’UTR
EGE-LJL-034-MSD-R2
----- LR ----
:CRISPR/Cas9
----- RR ----
 Targeted allele
EGE-LJL-034-MSD-F2
Tdtomato-2371F
3’UTR
 E2
GSG-3XFlag-P2A-Tdtomato
EGE-LJL-034-MSD-R2
Tdtomato-1R

### Slide 19
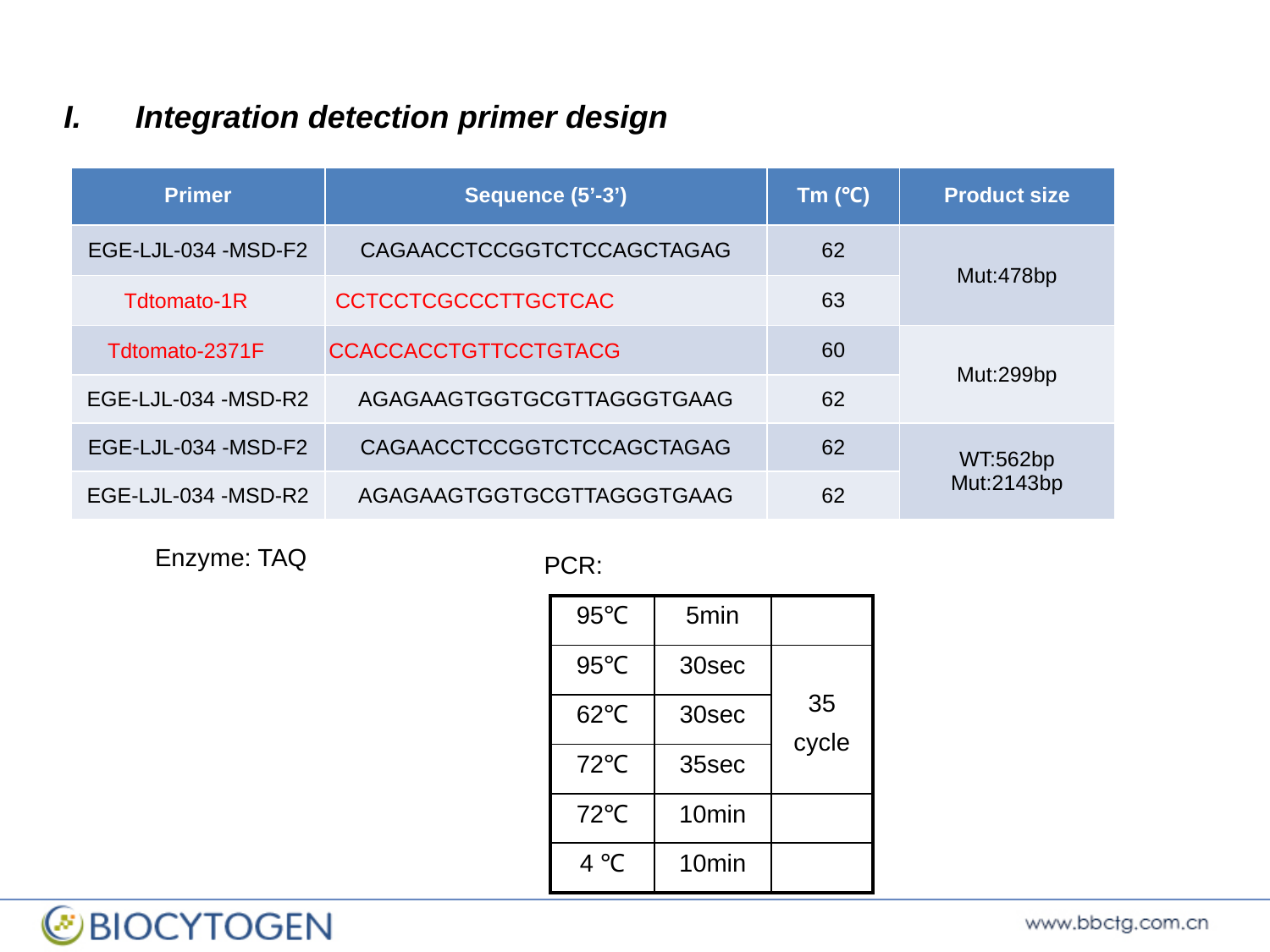

Integration detection primer design
| Primer | Sequence (5’-3’) | Tm (℃) | Product size |
| --- | --- | --- | --- |
| EGE-LJL-034 -MSD-F2 | CAGAACCTCCGGTCTCCAGCTAGAG | 62 | Mut:478bp |
| Tdtomato-1R | CCTCCTCGCCCTTGCTCAC | 63 | |
| Tdtomato-2371F | CCACCACCTGTTCCTGTACG | 60 | Mut:299bp |
| EGE-LJL-034 -MSD-R2 | AGAGAAGTGGTGCGTTAGGGTGAAG | 62 | |
| EGE-LJL-034 -MSD-F2 | CAGAACCTCCGGTCTCCAGCTAGAG | 62 | WT:562bp Mut:2143bp |
| EGE-LJL-034 -MSD-R2 | AGAGAAGTGGTGCGTTAGGGTGAAG | 62 | |
Enzyme: TAQ
PCR:
| 95℃ | 5min | |
| --- | --- | --- |
| 95℃ | 30sec | 35 cycle |
| 62℃ | 30sec | |
| 72℃ | 35sec | |
| 72℃ | 10min | |
| 4 ℃ | 10min | |

### Slide 20
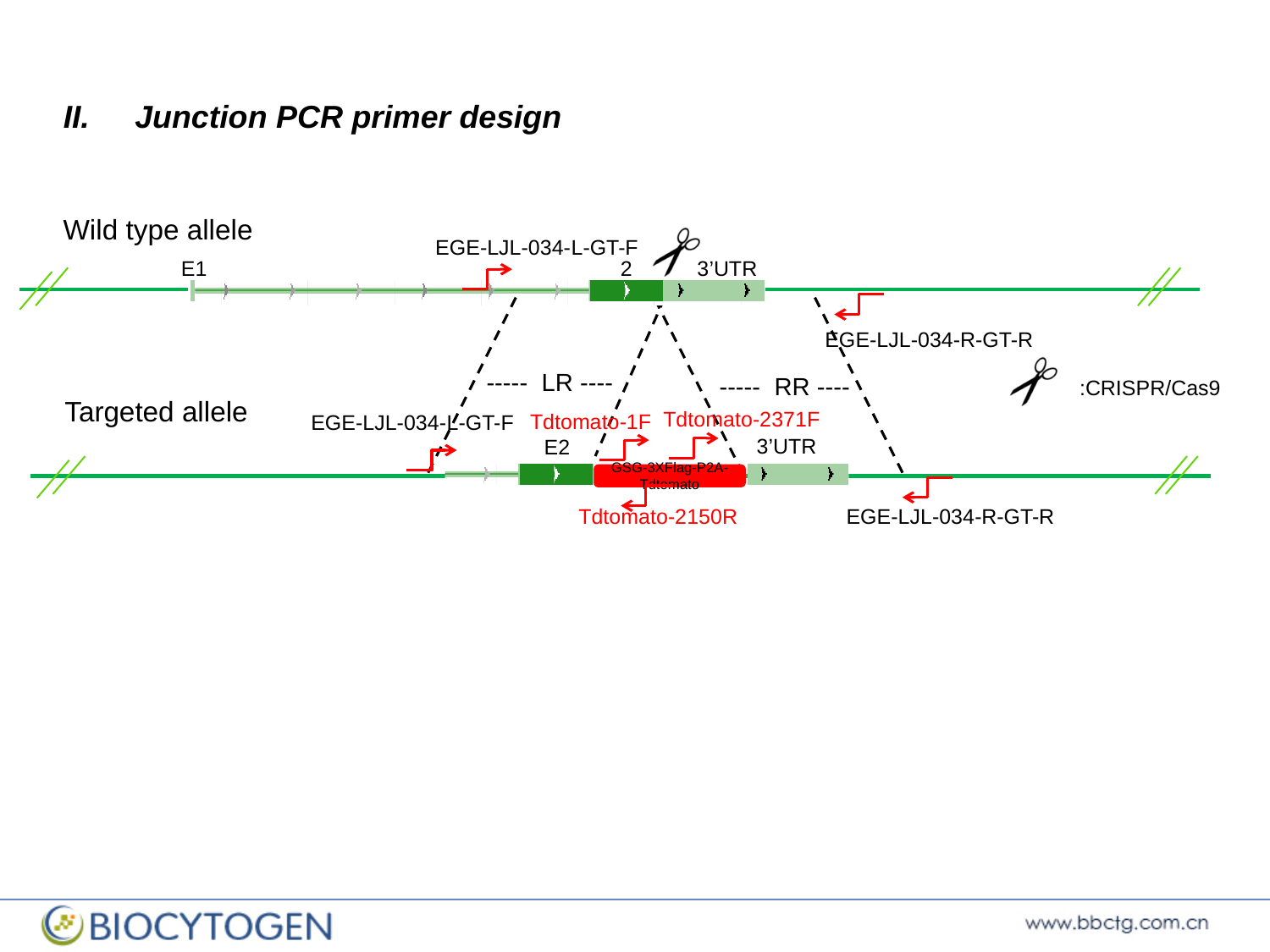

Junction PCR primer design
Wild type allele
EGE-LJL-034-L-GT-F
E1 2 3’UTR
EGE-LJL-034-R-GT-R
----- LR ----
:CRISPR/Cas9
----- RR ----
 Targeted allele
Tdtomato-2371F
Tdtomato-1F
EGE-LJL-034-L-GT-F
3’UTR
 E2
GSG-3XFlag-P2A-Tdtomato
Tdtomato-2150R
EGE-LJL-034-R-GT-R

### Slide 21
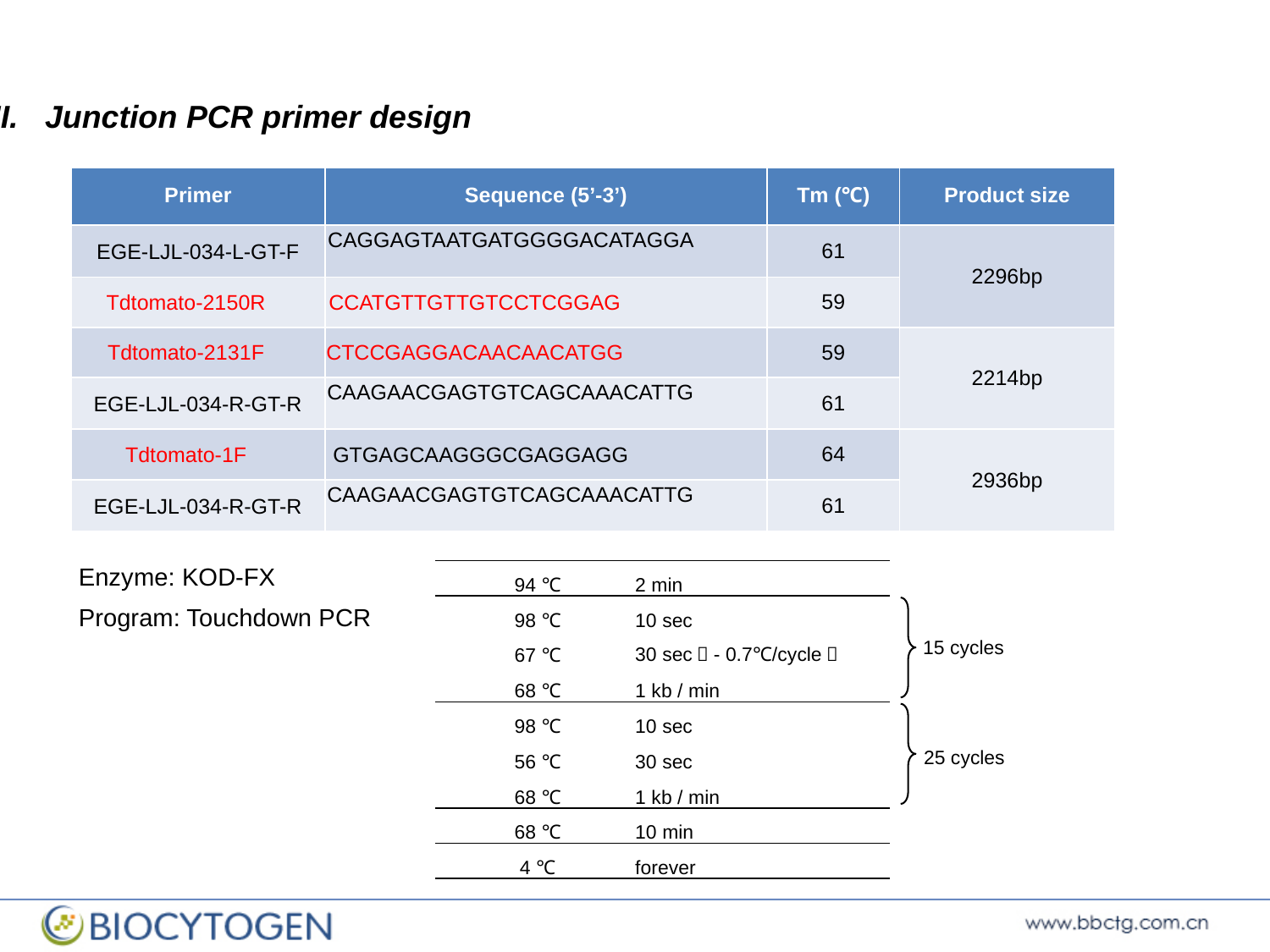

II. Junction PCR primer design
| Primer | Sequence (5’-3’) | Tm (℃) | Product size |
| --- | --- | --- | --- |
| EGE-LJL-034-L-GT-F | CAGGAGTAATGATGGGGACATAGGA | 61 | 2296bp |
| Tdtomato-2150R | CCATGTTGTTGTCCTCGGAG | 59 | |
| Tdtomato-2131F | CTCCGAGGACAACAACATGG | 59 | 2214bp |
| EGE-LJL-034-R-GT-R | CAAGAACGAGTGTCAGCAAACATTG | 61 | |
| Tdtomato-1F | GTGAGCAAGGGCGAGGAGG | 64 | 2936bp |
| EGE-LJL-034-R-GT-R | CAAGAACGAGTGTCAGCAAACATTG | 61 | |
Enzyme: KOD-FX
Program: Touchdown PCR
| 94 ℃ | 2 min |
| --- | --- |
| 98 ℃ | 10 sec |
| 67 ℃ | 30 sec（- 0.7℃/cycle） |
| 68 ℃ | 1 kb / min |
| 98 ℃ | 10 sec |
| 56 ℃ | 30 sec |
| 68 ℃ | 1 kb / min |
| 68 ℃ | 10 min |
| 4 ℃ | forever |
15 cycles
25 cycles

### Slide 22
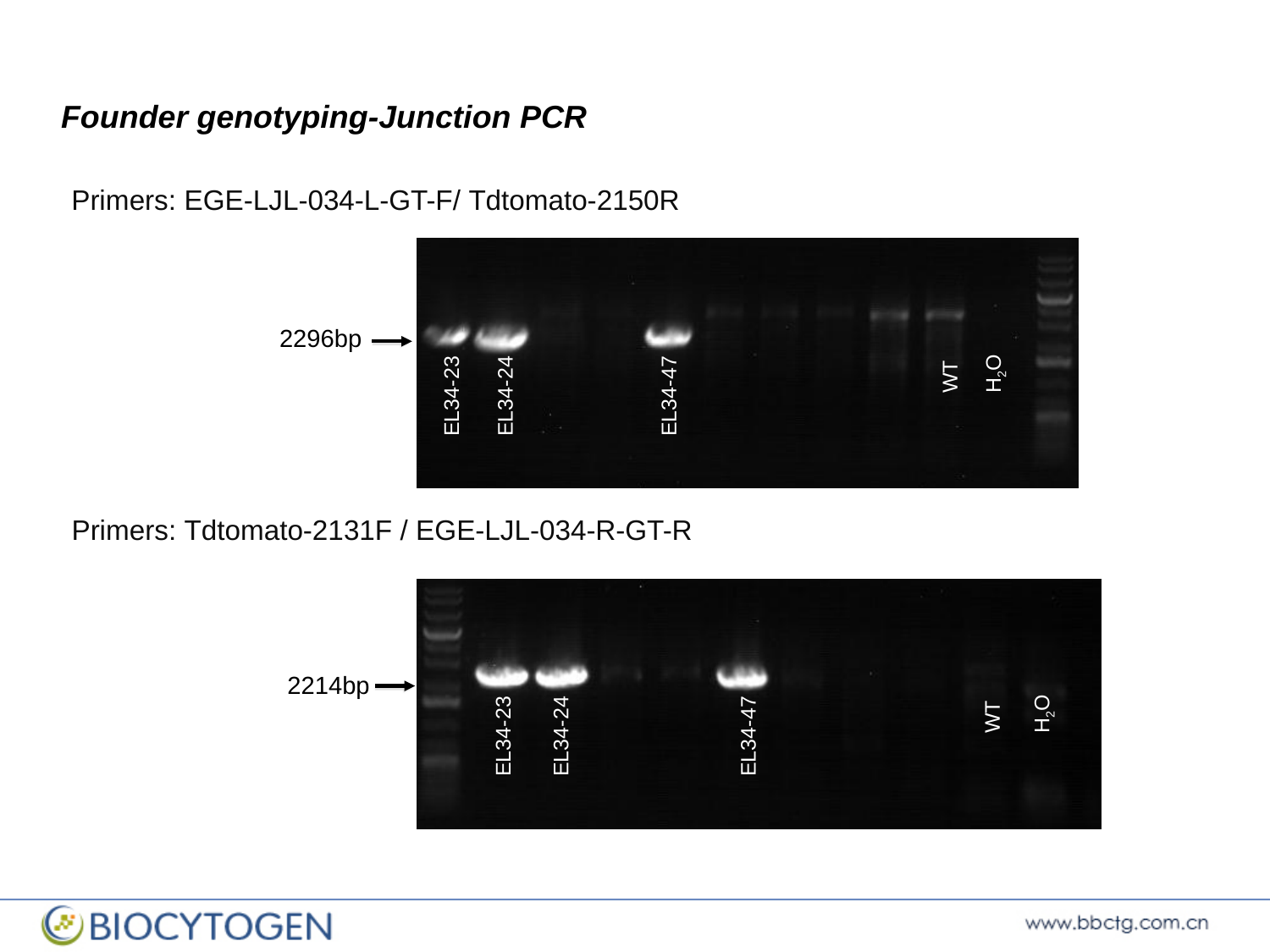

Founder genotyping-Junction PCR
Primers: EGE-LJL-034-L-GT-F/ Tdtomato-2150R
2296bp
EL34-47
WT
H_2_O
EL34-23
EL34-24
Primers: Tdtomato-2131F / EGE-LJL-034-R-GT-R
EE19-18
EE19-4
WT
H_2_O
2214bp
EL34-47
WT
H_2_O
EL34-23
EL34-24

### Slide 23
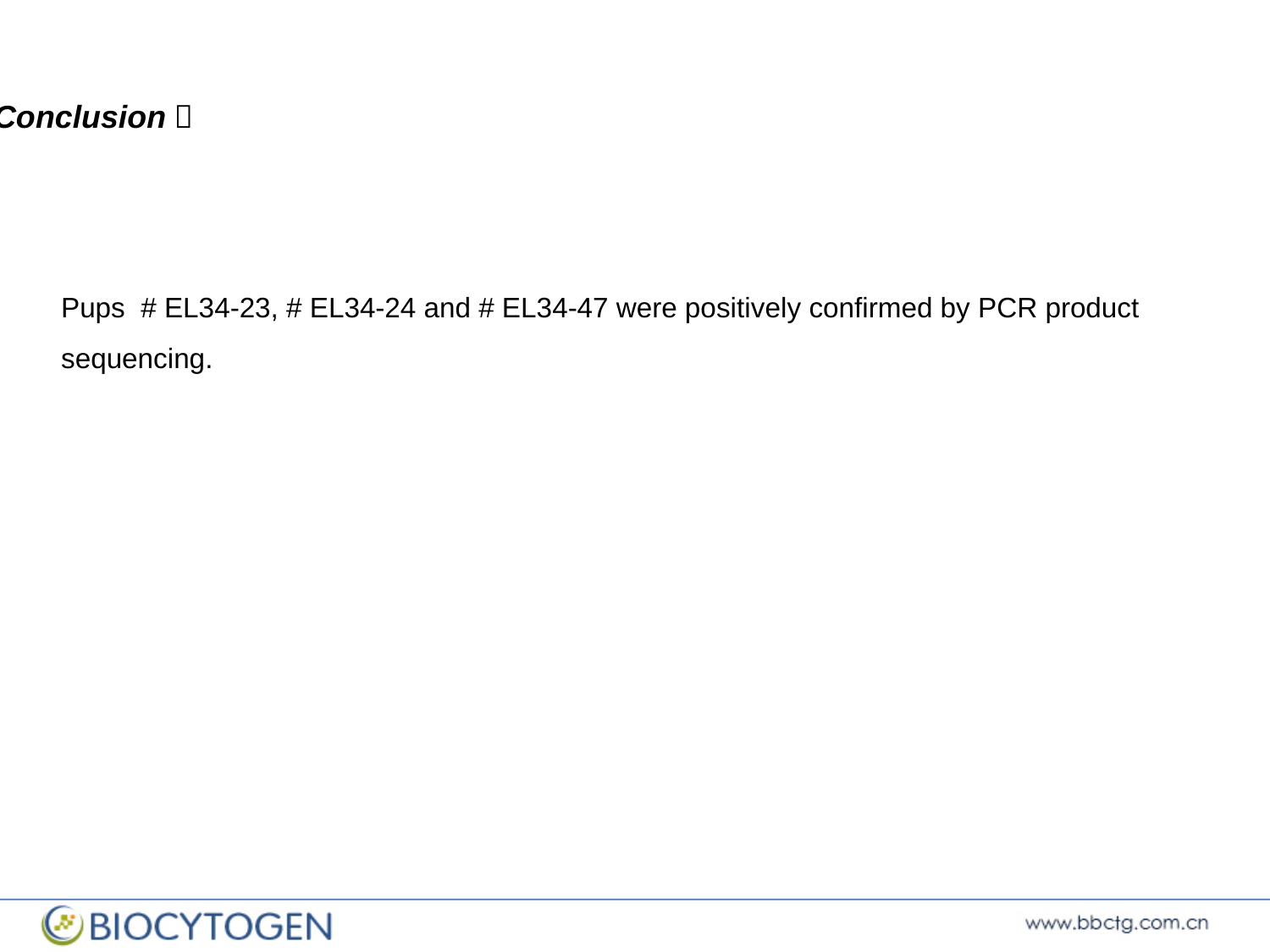

Conclusion：
Pups # EL34-23, # EL34-24 and # EL34-47 were positively confirmed by PCR product sequencing.
