## Supplementary File 2 for "A Novel *Tmem119-tdTomato* Reporter Mouse Model for Studying Microglia in the Central Nervous System"

#### 2.1.1 Primer design strategy

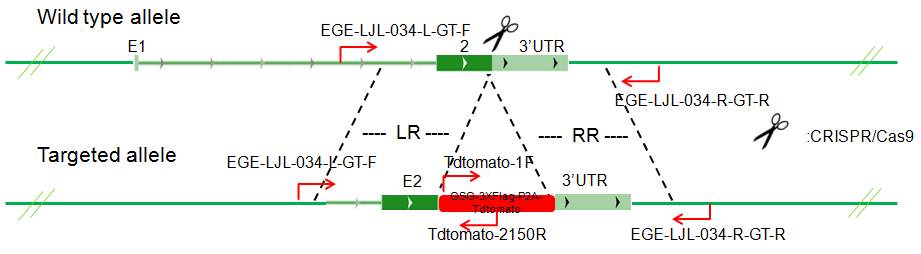

Figure 1: Schematic representation of the wild type and mutant/targeted alleles. Red arrows illustrate the primer’s localization. LR: left homologous arm; RR: right homologous arm.

#### 2.1.2 Primer sequences and predicted product sizes

| **Primer** | **Sequence (5’-3’)** | **Tm (℃)** | **Product size (bp)** |
| --- | --- | --- | --- |
| EGE-LJL-034-L-GT-F | CAGGAGTAATGATGGGGACATAGGA | 61 | Mut: 2296 |
| Tdtomato-2150R | CCATGTTGTTGTCCTCGGAG | 59 |  |
| Tdtomato-1F | GTGAGCAAGGGCGAGGAGG | 64 | Mut: 2936 |
| EGE-LJL-034-R-GT-R | CAAGAACGAGTGTCAGCAAACATTG | 61 |  |

Table 2: Primer set for detection of the knock in sequences integration into mice’s genome. Mut: representing the mutant/targeted allele.

#### 2.1.3 PCR reaction system

| **Reaction component** | **Volume (µl)** | **Final concentration** |
| --- | --- | --- |
| ddH_2_O | 1.9 | — |
| 2×KOD FX buffer | 10 | 1× |
| 2 mM dNTPs | 4 | 0.4 mM each |
| 10 μM Primer-F | 0.6 | 0.3 μM |
| 10 μM Primer-R | 0.6 | 0.3 μM |
| DMSO | 1 | 5% |
| 1U/μl KOD FX DNA Polymerase | 0.4 | 0.02 U/μl |
| 100-200 ng/20μl Template DNA | 1.5 | 5-10* ng/μl |

Table 3: *Final concentration of template DNA could be in a range of 5-10ng/μl.

#### 2.1.4 PCR reaction conditions

| **Step** | **Temp.** | **Time** | **Cycles** |
| --- | --- | --- | --- |
| 1 | 94 ℃ | 2 min | 1 |
| 2 | 98 ℃ | 10 sec | 15 |
| 3 | 67 ℃ | 30 sec（- 0.7℃/cycle） |  |
| 4 | 68 ℃ | 1 kb/min |  |
| 5 | 98 ℃ | 10 sec | 25 |
| 6 | 56 ℃ | 30 sec |  |
| 7 | 68 ℃ | 1 kb/min |  |
| 8 | 68 ℃ | 10 min | 1 |
| 9 | 4 ℃ | forever | 1 |

Table 4: Touch-down PCR program is applied for genotyping.

#### 2.1.5 Gel image

2.1.5.1 Primer set: EGE-LJL-034-L-GT-F/Tdtomato-2150R

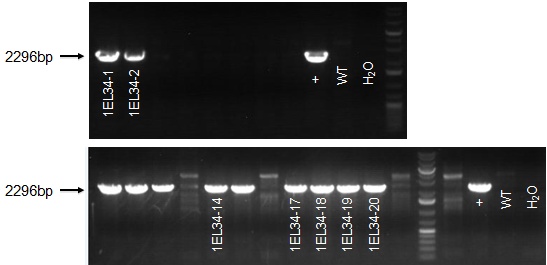

Figure 2: Gel image of the PCR products. DNA is separated by gel electrophoresis on a 1% agarose gel.

2.1.5.2 Primer set: Tdtomato-1F/EGE-LJL-034-R-GT-R

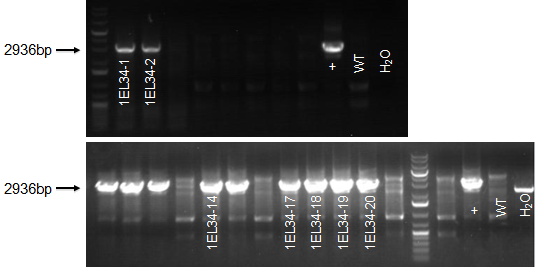

Figure 3: Gel image of the PCR products. DNA is separated by gel electrophoresis on a 1% agarose gel.

### 2.2 Sequencing analysis

Long fragment PCR products are sequenced to confirm the integration site of the knockin sequences in genome and no other mutation introduced in the recombination process. Sequencing results are blasted with the sequence of mutant/targeted allele which we designed. All the sequences of the PCR product are correct.

### 2.3 Short fragment PCR Screening

Germ line transmission of genetic recombination in F1 animal has been confirmed with long fragment PCR and sequencing analysis. Because of the product sizes of long fragment PCR are sometimes too long to be amplified. Short fragment PCR could be used conveniently for genotyping of all the progeny of F1 animals. Primers are designed for detection of the wild type allele and the mutation/targeted allele. So the genotype of all animals could be identified as homozygous, heterozygous or wild type.

#### 2.3.1 Primer design strategy

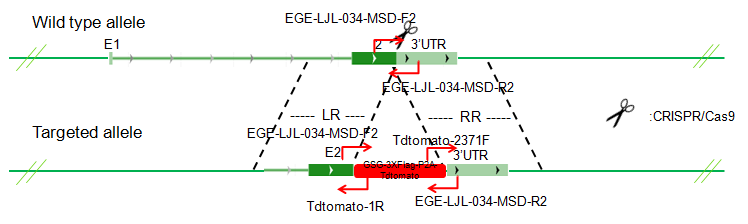

Figure 4: Schematic representation of wild type and mutant/targeted alleles. Red arrows illustrate the primer’s localization. LR: left homologous arm; RR: right homologous arm.

#### 2.3.2 Primer sequences and predicted product sizes

| **Primer** | **Sequence (5’-3’)** | **Tm (℃)** | **Product size (bp)** |
| --- | --- | --- | --- |
| EGE-LJL-034 -MSD-F2 | CAGAACCTCCGGTCTCCAGCTAGAG | 62 | Mut:478 |
| Tdtomato-1R | CCTCCTCGCCCTTGCTCAC | 63 |  |
| Tdtomato-2371F | CCACCACCTGTTCCTGTACG | 60 | Mut:299 |
| EGE-LJL-034 -MSD-R2 | AGAGAAGTGGTGCGTTAGGGTGAAG | 62 |  |
| EGE-LJL-034 -MSD-F2 | CAGAACCTCCGGTCTCCAGCTAGAG | 62 | WT:562  Mut:2143 |
| EGE-LJL-034 -MSD-R2 | AGAGAAGTGGTGCGTTAGGGTGAAG | 62 |  |

Table 5: Primer set and predicted product sizes for detection of the knock in sequences integration into genome. WT: wild type allele; Mut: mutant allele.

#### 2.3.3 PCR reaction system

| **Reaction component** | **Volume (µl)** | **Final concentration** |
| --- | --- | --- |
| ddH_2_O | 14 | — |
| 10×Taq buffer | 2 | 1× |
| 10 mM dNTPs | 0.5 | 250 μM each |
| 10 μM Primer-F | 0.5 | 0.25 μM |
| 10 μM Primer-R | 0.5 | 0.25 μM |
| 2.5 U/μl Taq DNA Polymerase | 0.5 | 0.0625 U/μl |
| 100-200 ng/μl Template DNA | 2 | 5-10* ng/μl |

Table 6: Taq DNA Polymerase reaction system. *Final concentration of template DNA could be in a range of 5-10ng/μl.

#### 2.3.4 PCR reaction conditions

| **Step** | **Temp.** | **Time** | **Cycles** |
| --- | --- | --- | --- |
| 1 | 95℃ | 3 min | 1 |
| 2 | 95℃ | 30 sec | 35 |
| 3 | 62℃ | 30 sec |  |
| 4 | 72℃ | 1 kb/min |  |
| 5 | 72℃ | 10 min | 1 |
| 6 | 4 ℃ | forever | 1 |

Table 7: Taq DNA Polymerase PCR program is applied for genotyping.

#### 2.3.5 Gel image

2.3.5.1 Primer set: EGE-LJL-034-MSD-F2/ Tdtomato-1R

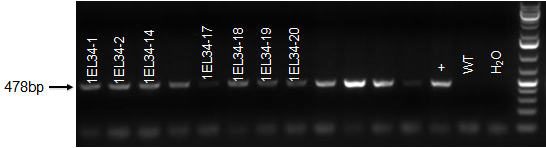

Figure 5: Gel image of the PCR products. DNA is separated by gel electrophoresis on a 1% agarose gel.

2.3.5.2 Primer set: Tdtomato-2371F/ EGE-LJL-034-MSD-R2

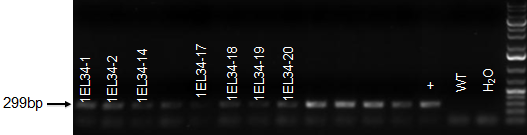

Figure 6: Gel image of the PCR products. DNA is separated by gel electrophoresis on a 1% agarose gel.

2.3.5.3 Primer set: EGE-LJL-034-MSD-F2/ EGE-LJL-034-MSD-R2

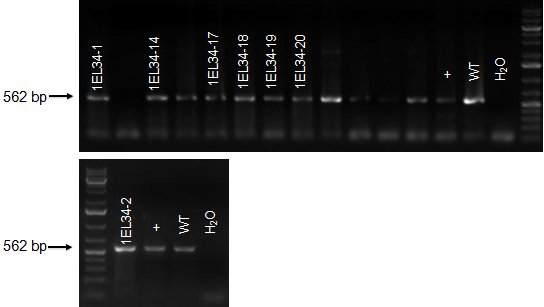

Figure 7: Gel image of the PCR products. DNA is separated by gel electrophoresis on a 1% agarose gel.

### 2.4 Determination of genotype from gel image results

| **EGE-LJL-034-MSD-F2/ Tdtomato-1R** | **Tdtomato-2371F/ EGE-LJL-034-MSD-R2** | **EGE-LJL-034-MSD-F2/ EGE-LJL-034-MSD-R2** | | **Genotype** |
| --- | --- | --- | --- | --- |
| **Mut** | **Mut** | **WT** | **Mut** |  |
| Y^1^ | Y | N^2^ | N ^3^ | Mut/Mut |
| Y | Y | Y | N | Mut/+ |
| N | N | Y | N | +/+ |

Table 8: Identification of three genotypes according to the expected products of short fragment PCR. Y^1^：Expected PCR product detected with gel electrophoresis. N^2^: No expected PCR product detected with gel electrophoresis. N^3^: Theoretically, when the primer set EGE-LJL-034-MSD-F2/ EGE-LJL-034-MSD-R2 is used for PCR, the product of mutation allele could be amplified. But sometimes this PCR product is too long to be amplified. So the result here is labeled as “N”. Mut/Mut: representing homozygous. Mut/+: representing heterozygous. +/+: representing wild type.
